# Entropic pressure controls oligomerization of *Vibrio cholerae* ParD2 antitoxin

**DOI:** 10.1101/2021.03.09.434581

**Authors:** Gabriela Garcia-Rodriguez, Yana Girardin, Alexander N. Volkov, Ranjan Kumar Singh, Gopinath Muruganandam, Jeroen Van Dyck, Frank Sobott, Wim Versées, Remy Loris

**Affiliations:** Structural Biology Brussels, Vrije Universiteit Brussel, Pleinlaan 2, Brussel, 1050, Belgium; VIB-VUB Center for Structural Biology, Vlaams Instituut voor Biotechnologie, Pleinlaan 2, Brussel, 1050, Belgium; Jean Jeener NMR Center, Vrije Universiteit Brussel, Pleinlaan 2, Brussel, 1050, Belgium; Department of Chemistry, Universiteit Antwerpen, Groenenborgerlaan 171, Antwerpen, 2020, Belgium; Astbury Centre for Structural Molecular Biology, University of Leeds, Leeds, LS29JT, United Kingdom

**Keywords:** Toxin-antitoxin system, Intrinsically disordered protein, Protein structure, Protein oligomerization, Molecular biophysics

## Abstract

ParD2 is the antitoxin component of the parDE2 toxin-antitoxin module from Vibrio cholerae and consists of an ordered DNA binding domain followed by an intrinsically disordered ParE-neutralizing domain. In absence of the C-terminal IDP domain, VcParD2 crystallizes as a doughnut-shaped hexadecamer formed by the association of eight dimers. This assembly is stabilized via hydrogen bonds and salt bridges rather than hydrophobic contacts. In solution, oligomerization of the full-length protein is restricted to a stable, open 10-mer or 12-mer, likely as a consequence of entropic pressure from the IDP tails. The relative positioning of successive VcParD2 dimers mimics the arrangement of *Streptococcus agalactiae* CopG dimers on their operator and allows for an extended operator to wrap around the VcParD2 oligomer.

## 1. Introduction

Like all organisms, bacteria have to deal with various kinds of stress that threaten their survival. This includes nutrient starvation, chemical stress arising from compounds including antibiotics or redox stress and physical stress such as heat or salt. To deal with this, they have developed different strategies, including the generation of persister cells, a specific stochastically induced dormant metabolic state that allows a subpopulation to survive antibiotics for which they are not resistant. Among the potential players in the bacterial stress response network are sets of small two-gene operons encoding a toxic protein and a corresponding neutralizing protein or RNA, known as toxin-antitoxin (TA) modules. They are often abundant in free-living bacteria and opportunistic pathogens (Pandey & Gerdes, 2005). For example, the well-known *Mycobacterium tuberculosis* contains at least 88 such modules while the closely related *M. smegmatis* only contains 5 (Shao et al., 2011).

The physiological role of toxin-antitoxin modules is highly debated. One of their potential functions is stabilization of mobile genetic elements (Gerdes et al., 1986; Szekeres et al., 2007). After initially been observed as stabilizing elements on low copy number plasmids, TA modules were also found on chromosomal regions that do not encode essential genes including integrative and conjugative elements, the superintegrons of *V. cholerae* and *V. vulnificus*, chromosome II, cryptic prophages and genomic and pathogenicity islands (Díaz-Orejas et al., 2017; Yao et al., 2018).

The most popular proposed function for TA modules is stress response (Gerdes, 2000; Gerdes et al., 2005; Hõrak & Tamman, 2017). Indeed, the activity of TA toxins is upregulated during episodes of stress. This was initially proposed to be a consequence of protease-dependent degradation of antitoxins, a mechanism that was recently challenged (LeRoux et al., 2020; Song & Wood, 2020a). It remains further unclear whether stress-related TA activation is an integral part of the stress response network, or is rather a side effect of the SOS response. Equally, the supposed role of TA modules in the onset of persistence remains unclear (Ronneau & Helaine, 2019).

Another function attributed to TA modules is protection against bacteriophages via abortive infection (Fineran, 2019; Lopatina et al., 2020; Song & Wood, 2020b). Last but not least, it should be considered that TA modules might be mere selfish genetic elements that in a number of cases adapted and have been conserved because they provide some additional beneficial properties such as the functions described above.

Toxin-antitoxin modules are divided into eight classes based on the nature of the antitoxin (protein or RNA) and the mechanism by which it counteracts the toxin (Page & Peti, 2016; Song & Wood, 2020a). The most common and best studied class of toxin-antitoxin modules are the type II modules where that antitoxin encodes a protein that counteracts the toxin via the formation of a tight non-covalent complex. Type II modules can be further classified in an increasing number of non-related families. Among those, the *parDE* family of TA modules, although one of the early families that were discovered, is relatively understudied (Roberts & Helinski, 1992). For two members, *parDE* from *Escherichia coli* plasmid RK2 and *parDE2* from *Vibrio cholerae*, the target of the ParE toxin was identified as gyrase (Jiang et al., 2002; Yuan et al., 2010). Like the more intensively studied CcdB, ParE poisons gyrase by stabilizing the so-called cleavable complex between Gyrase and DNA. This leads to double strand breaks, activation of the SOS response and ultimately, if the ParE toxin is not counteracted by the antitoxin ParD, cell death.

Currently the Protein data bank contains an NMR structure for ParD from plasmid RK2, an NMR structure of *E. coli* PaaA2, which is a truncated ParD protein lacking a DNA binding domain, and crystal structures for ParD-ParE complexes from *Caulobacter crescentus* and *Mesorhizobium opportunistum*, and from the *parDE*-like *paaR2-paaA2-parE2* from *E. coli* O157:H7 (Oberer et al., 2007; Sterckx et al., 2014; Dalton & Crosson, 2010; Aakre et al., 2015; Sterckx et al., 2016). For neither of these ParE proteins the target has been identified and for *E. coli* ParE2, evidence was presented that it does not interact with the DNA Gyrase A subunit (Sterckx et al., 2016). RK2, *M. opportunistum* and *C. crescentus* ParD fold into an N-terminal ribbon-helix-helix DNA binding domain that is followed by a domain which for RK2 ParD is unfolded in solution and for *C. crescentus* and *M. opportunistum* ParD folds into a helix-helix-strand conformation that wraps around the ParE toxin. *E. coli* PaaA2 lacks the N-terminal DNA binding domain and is mostly disordered in solution but adopts the same conformation as the *C. crescentus* or *M. opportunistum* C-terminal domain when bound to its cognate ParE2.

The genome of *Vibrio cholerae* contains three *parDE* modules, all located in the superintegron on chromosome II (Yuan et al., 2011). The *parDE1* and *parDE3* modules have identical open reading frames and regulatory sequences while the ParD and ParE proteins encoded by the *parDE2* module share 14% and 22% sequence identity to their *parDE1/3* counterparts. ParE2 was shown to inhibit Gyrase *in vitro* and to bind to an epitope on the Gyrase A subunit that is different from the one that is targeted by F-plasmid CcdB (Yuan et al., 2010). *In vivo*, both ParE1/3 and ParE2 inhibit cell division, activate the SOS response and contribute to the degradation of chromosome I upon loss of chromosome II.

Transcription regulation of TA modules is often complex and involves the ratio-dependent interplay between toxin and antitoxin (Garcia-Pino et al., 2010; Jurėnas et al., 2019; Page & Peti, 2016; Vandevelde et al., 2017; Xue et al., 2020). In these mechanisms, intrinsically disordered segments on antitoxin or toxin often play a major role (De Jonge et al., 2009; Garcia-Pino et al., 2016; Loris & Garcia-Pino, 2014; Talavera et al, 2019). Little information is known, however on the regulation of *parDE* modules. Early work on plasmid RK2 *parDE* suggested that ParD alone can act as a repressor for the operon (Roberts et al., 1993), but it is not known whether the ParE toxin modulates this action. Here we describe the crystal structure of *Vibrio cholerae* ParD2 (*Vc*PardD2 hereafter) and show that inter-dimer contacts in the crystal mimic the functional arrangement of the structurally related *Streptococcus agalactiae* CopG bound to its DNA target (PDB entry 1B01 - Gomis-Ruth et al., 1998). While in the crystal a doughnut-shaped hexadecamer is observed, in solution smaller decamers and dodecamers are present that form a partial doughnut with otherwise similar inter-subunit contacts. The partial disruption of the full hexadecameric ring can possibly be attributed to steric pressure from the intrinsically disordered C-terminal tails that are absent in the crystallized entity.

## 2. Material and Methods

### 2.1 Expression and purification

The coding region of the *V. cholerae parDE2* operon was cloned into pET28a, placing a T7 promotor upstream of the *parD2* gene and adding a His-tag at the C-terminus of *Vc*ParE2. BL21 (DE3) *E. coli* cells were transformed with this vector and grown at 37°C in LB supplemented with 50 µg ml^-1^ kanamycin. Expression was induced at OD^600^ = 0.6 by adding 0.5 mM Isopropyl β-D-1-thiogalactopyranoside. The cultures were incubated overnight at 20°C, after which the cells were harvested by centrifugation at 5000 rpm, 4°C for 15 minutes.

The cells were re-suspended in 20 mM Tris pH 8.0, 500 mM NaCl, 2mM β-mercaptoethanol and broken with a cell cracker after adding 50 µg ml^-1^ DNase and 2 mM MgCl_2_. After centrifugation at 19500 rpm, 4°C for 35 minutes, the supernatant was filtered using a 0.45 µm filter and then loaded onto a 5 ml HisTrap™ HP nickel sepharose column (GE Healthcare) equilibrated with the same buffer. The column was washed with 20 mM Tris pH 8.0, 500 mM NaCl, 2mM β-mercaptoethanol and the *Vc*ParD2-*Vc*ParE2 complex was subsequently eluted using a step gradient of imidazole (0 – 10 - 25 - 50 - 250 - 500 mM) in the same buffer. The resulting samples were loaded onto a Superdex 200 16:90 SEC column (GE Healthcare) pre-equilibrated with 20 mM Tris pH 8.0, 500 mM NaCl, 2mM β-mercaptoethanol to obtain pure ParD2-ParE2 complex.

The *Vc*ParD2-*Vc*ParE2 complex was re-loaded on a Ni-NTA column followed by washing with 20 mM Tris pH 8.0, 0.5 M NaCl, 10 % ethylene glycol. *Vc*ParD2 was eluted by applying a step gradient of guanidinium hydrochloride (GndHCl - 0 - 2.5 - 5 M) in 20 mM Tris pH 8.0, 500 mM NaCl, 2 mM β-mercaptoethanol. The resulting *Vc*ParD2 fractions were pooled and dialyzed against 20 mM Tris pH 8.0 150 mM NaCl prior to a final SEC run on a Superdex 200 16:60 column (GE healthcare).

The Ni-NTA column was subsequently washed with (1) 20 mM Tris pH 8.0, 25 mM NaCl, 5 % glycerol; (2) 20 mM Tris pH 8.0, 150 mM NaCl, 5 % glycerol and (3) 20 mM Tris pH 8.0, 250 mM NaCl, 2 mM β-mercaptoethanol to refold *Vc*ParE2 on the column. *Vc*ParE2 was eluted using a step gradient of imidazole (25 - 50 - 250 - 500 mM) and further purified on a Superdex 75 16:60 column. Purity of all samples was checked using SDS-PAGE and nano-electrospray ionization-time of flight (nESI-TOF) mass spectrometry.

### 2.2 Crystallization, data collection and structure determination

*Vc*ParD2 in 20 mM Tris pH 8.0, 150 mM NaCl was concentrated up to 20 mg ml^-1^. For screening crystallization conditions, 0.1 µl of protein solution were mixed with 0.1 µl of reservoir solution in a sitting drop equilibrated against 100 µl of reservoir solution using a Mosquito HTS robot (http://ttplabtech.com/). Crystals were observed after approximately three months in 0.2 M Lithium sulfate, 0.1 M MES pH 6.0 and 20 % w/v PEG 4000. Crystals were transferred to precipitant solution supplemented with 25 % PEG 400 for cryo-protection and immediately flash-cooled in liquid nitrogen. Data were collected at Proxima 2A beamline of the Soleil Synchrotron facility (Gif-sur-Yvette, Paris, France). All data were indexed, integrated and scaled with XDS (Kabsch, 2010) via the XDSME interface (Legrand, 2017). Analysis of solvent content was performed using the CCP4 program MATTHEWS_COEF (Kantardjieff & Rupp, 2003).

The structure of *Vc*ParD2 was determined by molecular replacement with Phaser-MR (McCoy, 2007) using the dimer of the N-terminal domain (residues 1-50) of *C. crescentus Cc*ParD2 (PDB ID 3kxe, 63 % Seq. Id. - Dalton & Crosson, 2010). Four copies of this N-terminal dimer were placed in the asymmetric unit (AU). Several cycles of phenix refinement (Afonine et al., 2012) and manual model building in Coot (Emsley et al., 2010) improved the phases. Further iterative cycles in *phenix*.*refine* were performed using an intensities-based maximum likelihood target function including TLS and NCS refinement with automated group determination as implemented in *phenix*.*refine*. Strong NCS restraints were applied throughout the refinement, leading to a relatively high value for R_work_ but a comparably small difference between R_work_ and R_free_. We assumed because of the low resolution, that all eight chains are identical, and choose to include the same number of amino acids. Consequently, rather poor fits for residues 3-4 of some of the chains remain. Refinement statistics are given in Table I. Structural homologs where identified using DALI (http://ekhidna2.biocenter.helsinki.fi/dali - Holm, 2020). Macromolecular interfaces were analyzed using the PISA server (http://www.ebi.ac.uk/pdbe/prot_int/pistart.html - Krissinel & Henrick, 2007).

### 2.3. Circular dichroism spectroscopy

Protein concentrations were determined spectrophotometrically assuming extinction coefficients at 280 nm of 2980 M^-1^ cm^-1^ for *Vc*ParD2 and 15470 M^-1^ cm^-1^ for *Vc*ParE2-His as determined from the amino acid sequences using the Expasy.org ProtParam tool (Gasteiger et al., 2005). CD spectra were recorded at room temperature on a Jasco J-715 spectropolarimeter in 20 mM Tris pH 8.0, 150 mM NaCl, 1 mM TCEP and at a concentration of 0.15 mg ml^-1^.

### 2.4. Analytical SEC and SEC-MALS

Analytical SEC experiments were performed using Superdex Increase 200 10:300 and Superdex Increase 75 10:300 columns (GE Healthcare) equilibrated with 20 mM Tris pH 8, 500 mM NaCl, 1 mM TCEP. The BioRad gel filtration standards (bovine thyroglobulin, 670 kDa; bovine γ-globulin, 158 kDa, chicken ovalbumin, 44 kDa; horse myoglobin, 17 kDa; vitamin B12, 1.35 kDa) were used to make a standard curve with the logarithm of the molecular weight of the standards as a function of their elution volume.

Multi-angle light scattering experiments coupled to SEC (SEC-MALS) were performed using a HPLC system (Waters) connected in line with Treos II (Wyatt Technology) light scattering detector (using 3 angles) followed by a Shodex Refractive index detector (RI-501). The shodex-K402.5-4F SEC column was connected to the SEC-MALS system and equilibrated with 2-3 column volumes of the corresponding running buffer. A dilution series of *Vc*ParD2 samples was prepared in the same buffer starting from 18 mg ml^-1^ to 0.1 mg ml^-1^. For each dilution, 10 µl were injected. A BSA sample of 1 mg ml^-1^ was used as a standard for calibration. Data was processed, and consequently molar mass was determined using the Astra 7.1.4 software.

### 2.5. Native mass spectrometry

For native MS the ParD2 was transferred into different ammonium acetate concentrations (50, 150 and 500 mM) at different pH (8.0 and 5.6) using Biospin buffer exchange columns (Bio-Rad Laboratories N.V., Temse, Belgium). After buffer exchange, concentration of the protein in different buffer conditions was determined by NanoDrop P2000 spectrophotometer (Thermo Scientific, Waltham, Massachusetts, USA).

For each buffer condition, samples were introduced into the mass spectrometer at *Vc*ParD2 concentrations of 0.1, 1.0 and 2.5 mg ml^-1^. Nano-electrospray ionization (ESI) was performed using 3-4 μl of sample loaded into home-made gold-coated borosilicate glass capillaries, with spray voltages applied in the range +1.5-2.0 kV. The spectra were recorded on an ion mobility enabled time-of-flight mass spectrometer (Synapt G2 HDMS, Waters, Wilmslow, UK). To achieve gentle, native-like conditions, the instrument parameters were carefully optimized in order to avoid ion activation and preserve higher-order structure of *Vc*ParD2 in the MS. The values optimized for *Vc*ParD2 are the sampling cone [50 V], extraction cone [2 V], trap collision energy [20 V], trap DC bias [45 V], and transfer collision energy [4 V]. Pressure in the source region (backing) and in the trap cell (collision gas) were 5.5 10^−2^ and 3.1 10^−2^ mbar respectively. Spectra were externally calibrated using a 10 mg ml^-1^ solution of cesium iodide. Analyses of the acquired spectra were performed using Masslynx version 4.1 (Waters, Wilmslow, UK). Native MS spectra were smoothed (extent depending on size of the complexes) and additionally centered for calculating molecular weights.

### 2.6. Small angle X-ray scattering

SAXS data were collected at the SWING beamline (Soleil, Saint-Aubin, France) in HPLC mode using an Agilent BioSEC 3-300 column at a protein concentration of 12 mg ml^-1^ in 20 mM Tris, pH 8.0, 150 mM NaCl, 1 mM TCEP at 19 °C. Protein samples were briefly spun down before loading on the SEC column. 50 μl of *Vc*ParD2 was injected onto the column with a constant flow of 0.2 ml min^-1^. The final scattering curve (after buffer subtraction) was generated for each sample after a range of scattering curves around the peak (with equivalent *R*_*g*_ values) was normalized and averaged. The *R*_*g*_ values were derived from the Guinier approximation at small *q* values while the *I*_*o*_ parameter was estimated by extrapolation to *q* = 0 using the ATSAS suite (Franke et al., 2017). Molecular weights were determined by the Bayesian estimation implemented in Primus (ATSAS suite).

All simulations were performed in Xplor-NIH v 2.49 (Schwieters et al., 2018), starting from the 16-mer X-ray structure of the *Vc*ParD2 solved in this work. Residues not resolved in the X-ray structure, including the disordered C-terminal tail, were added in Xplor-NIH using standard topologies for individual amino acids, followed by minimization of the energy function consisting of the standard geometric (bonds, angles, dihedrals, and impropers) and steric (van der Waals) terms.

For the refinement against the experimental SAXS data, the positions of the structured protein regions were kept fixed, while the disordered protein termini (including residues 1-4 and 48-81) were given full degree of freedom. The computational protocol comprised an initial simulated annealing step followed by the side-chain energy minimization as described before (Schwieters & Clore, 2014). In addition to the standard geometric and steric terms, the energy function included a knowledge-based dihedral angle potential and the SAXS energy term incorporating the experimental data (Schwieters et al., 2014). Truncated SAXS curves (q < 0.5 Å^-1^) were used as the sole experimental input.

In each refinement run, 100 structures were calculated and 10 lowest-energy solutions – representing the best agreement with the experimental data – retained for the subsequent analysis. The agreement between the experimental and calculated SAXS curves (obtained with the calcSAXS helper program, which is part of the Xplor-NIH package) was assessed by calculating the χ^2^ as in equation 1.

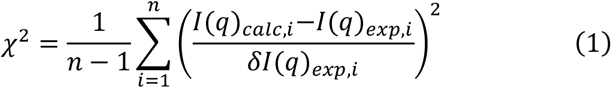

where *I*(*q*)_*calc,i*_ and *I*(*q*)_*exp,i*_ are the scattering intensities at a given *q* for the calculated and experimental SAXS curves, *δ I*(*q*)_*exp,i*_ is an experimental error on the corresponding *I*(*q*)_*exp,i*_ value, and *n* is the number of data points defining the experimental SAXS curve.

## 3. Results

### 3.1. Purification of *Vc*ParD2 and *Vc*ParE2

We initially attempted to express *Vc*ParD2 in *E. coli* from a pET28a expression vector that adds N-terminal His-tag followed by TEV cleavage site. These attempts led to very poor yields, possibly as a consequence of proteolytic degradation of *Vc*ParD2 in absence of *Vc*ParE2. We therefore altered our strategy and co-expressed *Vc*ParD2 and *Vc*ParE2 to obtain a *Vc*ParD2-*Vc*ParE2 complex. In order to obtain isolated *Vc*ParD2 and *Vc*ParE2 we used an on-column unfolding-refolding protocol similar to the ones developed previously for Phd/Doc, MazEF and HigBA (Sterckx *et al*., 2015). *Vc*ParE2 is His-tagged at its N-terminus and the *Vc*ParD2-*Vc*ParE2 complex is trapped in a Ni-NTA column. First, after extensive washing of the column, the *Vc*ParD2-*Vc*ParE2 complex is eluted and further cleaned up using SEC run on a Superdex 200 column. This step removes contaminating proteins of lower molecular weight that may otherwise co-migrate with *Vc*ParD2 or *Vc*ParE2 in further steps. Subsequently, the *Vc*ParD2-*Vc*ParE2 complex is re-loaded onto the Ni-NTA column. *Vc*ParD2 can then be eluted by applying a GndHCl gradient. This allows us to prepare *Vc*ParD2 in absence of an affinity tag or other cloning artefact, ensuring that the quaternary structure is not affected by the presence of a tag. After elution of *Vc*ParD2, the protein is refolded by dialysis. The refolded protein is then further polished by a SEC run on a Superdex 200 column (Supplementary Figure S1).

Our method has the additional advantage that also *Vc*ParE2 can be obtained, as like most other TA toxins, *Vc*ParE2 cannot be expressed in absence of its cognate antitoxin. The *Vc*ParE that remains trapped is refolded on the Ni-NTA column in a three-step procedure that consists of subsequent washing with (1) 20 mM Tris pH 8.0, 25 mM NaCl, 5 % glycerol, (2) mM Tris pH 8.0, 150 mM NaCl, 5 % glycerol and (3) 20 mM Tris pH 8.0, 250 mM NaCl, 2 mM β-mercaptoethanol. *Vc*ParE2 is subsequently eluted using a step gradient of imidazole. The resulting protein is then further applied onto a Superdex 75 column to obtain a sample that is pure on SDS-PAGE.

Both *Vc*ParE2 and *Vc*ParD2 show a CD spectra that are compatible with folded proteins (Figure 1a). *Vc*ParD2 is almost exclusively composed of α-helix while *Vc*ParE2 contains both α-helix and β-sheet. The SEC profile of *Vc*ParE2 is compatible with a globular monomer (Figure 1b). *Vc*ParD2, nevertheless, elutes at an apparent molecular weight of 170 kDa that is much higher than 18 kDa that is expected for a dimer (Figure 1c). While the presence of an IDP region will lead to aberrant migration in SEC, the deviation seems too large to be solely explainable by this phenomenon and a higher oligomer is likely formed. Both elution profiles are nevertheless indicative of monodisperse samples and together with the CD spectra suggest correctly folded proteins.

**Figure 1.**
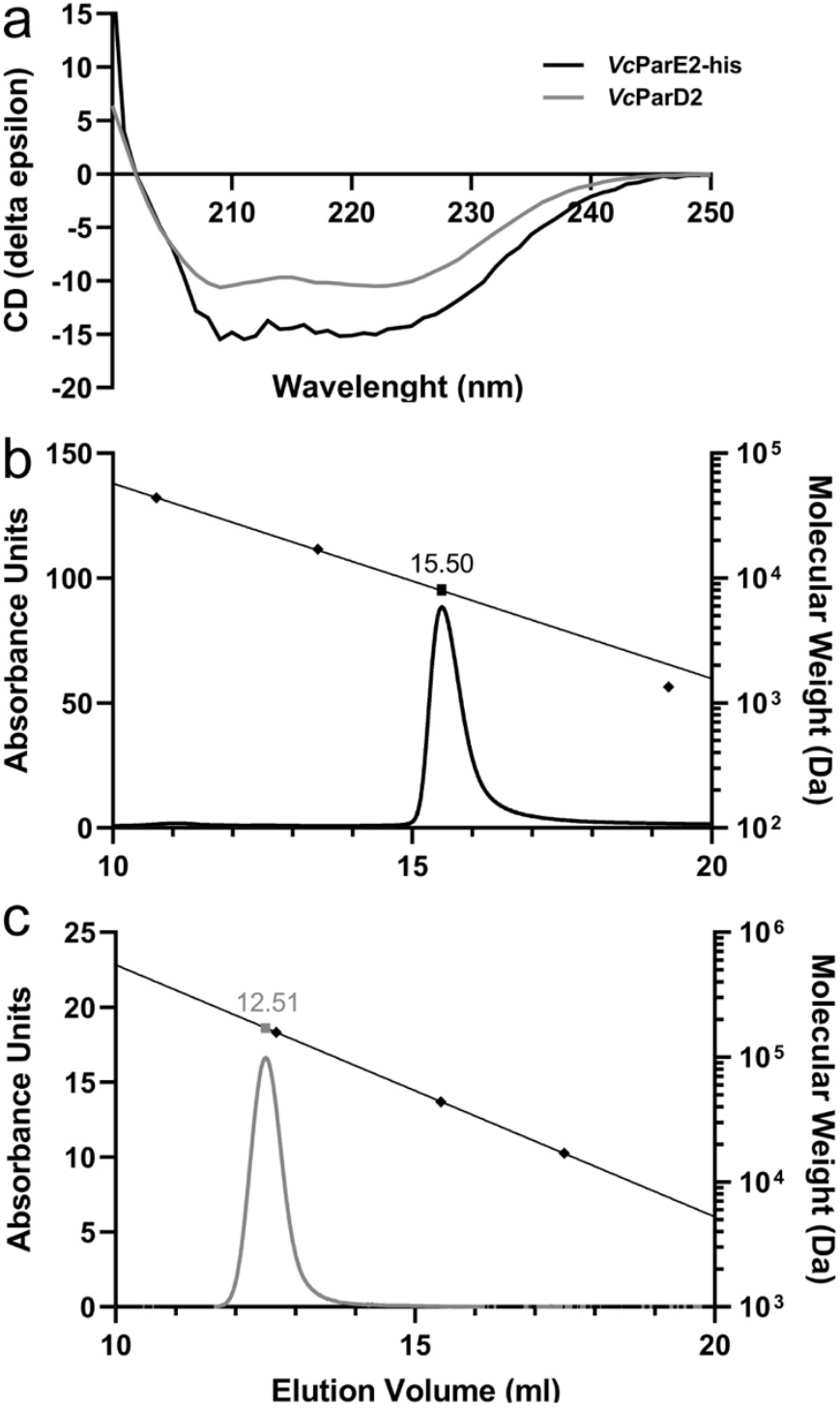
*Vc*ParE2 and *Vc*ParD2 are monodisperse, folded proteins. **a**. CD spectra of 0.15 mg ml^-1^ *Vc*ParD2 (grey) and *Vc*ParE2-His (black) in 20 mM Tris pH.0 8, 150 mM NaCl, 1 mM TCEP. **b**. Analytical SEC elution profile of 0.15 mg ml^-1^ *Vc*ParE2 in 20 mM Tris pH 8.0, 150 mM NaCl, 1 mM TCEP on a Superdex Increase 75 10:300 column. The elution volumes of the molecular weight standards (bovine γ-globulin 158 kDa; chicken ovalbumin 44. kDa; horse myoglobin 17 kDa; Vitamin B12 1.35 kDa) are shown as diamonds together with a linear regression (black line). **c**. Analytical SEC elution profile of 0.1 mg ml^-1^ *Vc*ParD2 in 20 mM Tris pH 8.0, 150 mM NaCl, 1 mM TCEP on a Superdex Increase 200 10:300 column. The elution volumes of the molecular weight standards (bovine thyroglobulin 670 kDa; bovine γ-globulin 158 kDa; chicken ovalbumin 44 kDa) are shown as diamonds together with a linear regression (black line). Each figure caption should be a single paragraph (style: IUCr figure caption; this style applies the figure numbering).

### 3.2. The *Vc*ParD2 monomer and dimer

The *Vc*ParD2 sample formed crystals that diffract to 3.1 Å resolution only after 3 months. The structure was determined by molecular replacement using the dimer of the N-terminal domain of *Caulobacter crescentus* ParD2 (*Cc*ParD2) (Dalton & Crosson, 2010), which shows 63% sequence identity to *Vc*ParD2 (54% for the complete protein chain), as search model. The structure was refined to an R_work_ of 0.2709 and an R_free_ of 0.2988 (Table I). The asymmetric unit contains four dimers of the N-terminal domain of *Vc*ParD2 and the corresponding models encompass residues Lys3 to Arg51. The N-terminal 2 and C-terminal 32 residues are not visible in the electron density map.

Although we did not succeed to determine the length of the polypeptide present in the crystal, it is highly unlikely that 8 intact *Vc*ParD2 chains are present in the asymmetric unit. The solvent contents calculated based on the visible part of the *Vc*ParD2 chains is 53%, a very reasonable value. In case of intact 81 amino acid *Vc*ParD2 chains, this would reduce to 22%. Although such a low solvent contents is not entirely impossible, the estimated probability is below 1% and in such a case the crowded environment would be expected to induce structure in the IDP region as is observed for *Mycobacterium tuberculosis* YefM (Kumar et al., 2008) and in part for bacteriophage P1 Phd (Garcia-Pino et al., 2010). Furthermore, degradation of IDP regions of antitoxins during crystallization has been observed before (Hadži et al., 2017).

As expected, the *Vc*ParD2 N-terminal domain (*Vc*ParD2_N_) adopts a ribbon-helix-helix DNA binding fold (Figure 2a), with a Cα RMSD of 0.7 Å for 47residues equivalent to *Cc*ParD2_N_ (Dalton & Crosson, 2010 - PDB entry 3kxe). Its topology consists of four helices in an open array of two hairpins often found in bacterial and phage repressors. From a DALI search, a number of additional structurally related transcription factors are identified that not only include the expected N-terminal domains of *Mesorhizobium opportunistum* ParD3, but also CopA, the antitoxin from the newly identified type II TA module ParE_SO_/CopA_SO_ from *Shewanella oneidensis*, the N-terminal domain of antitoxin AtaR from the *E. coli* AtaT-AtaR TA pair, the non-TA related 45-residue long transcriptional repressor CopG from *Streptococcus agalactiae* as well as the Arc repressor from *Salmonella* bacteriophage P22 (Table II and Supplementary Figure S2). Plasmid RK2 ParD, with only 8% sequence identity, shows up only as a low-ranking hit in this search.

**Figure 2.**
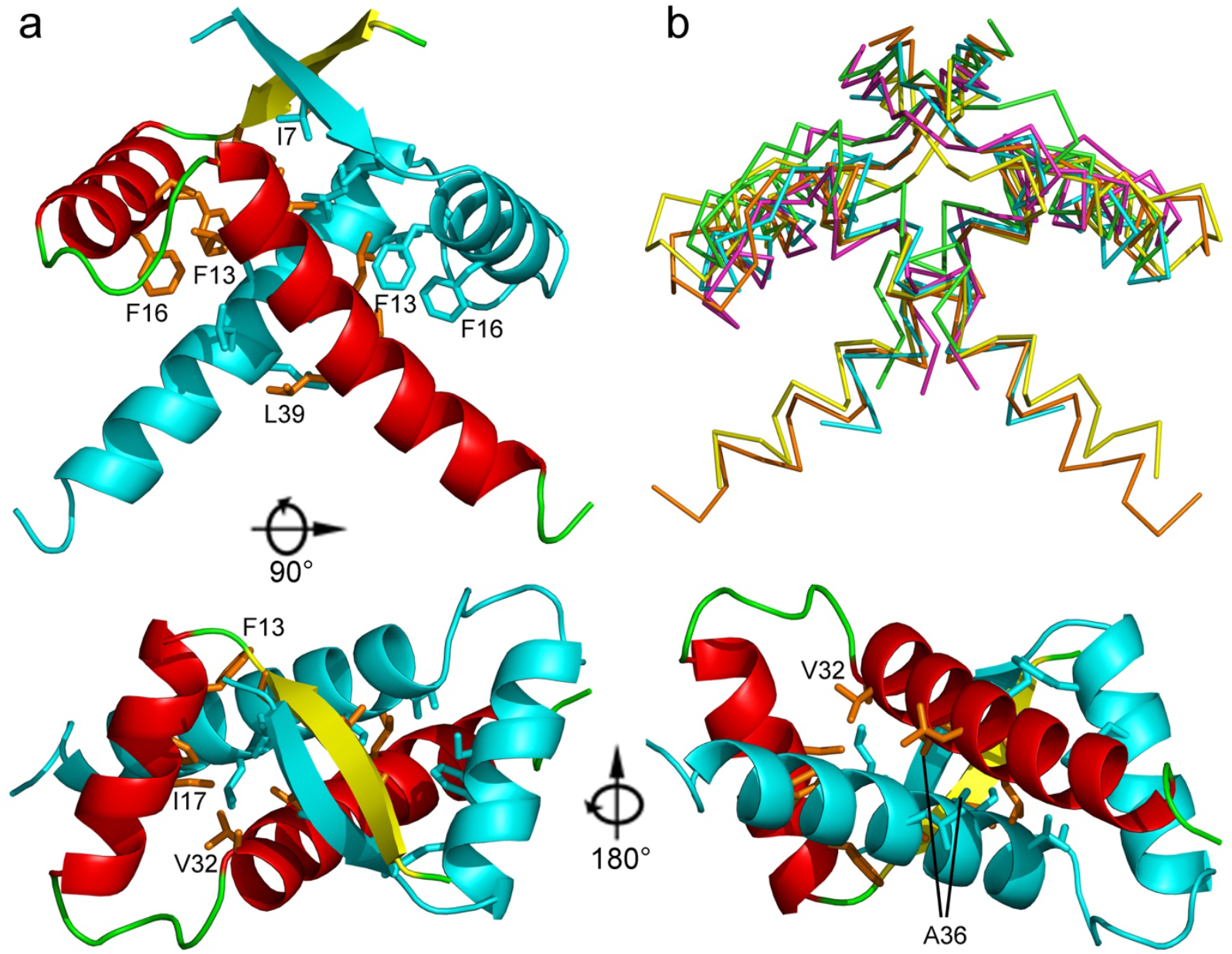
Crystal structure of *Vc*ParD2. **a**. Cartoon representation of the *Vc*ParD2 dimer (residues 3-51) in three orientations. One monomer is colored according to secondary structure (β-strand yellow, α-helix red and loop regions green). The second monomer is colored cyan. **b**. Superposition of the *Vc*ParD2_N_ dimer (orange) with the dimers of other RHH-type transcription factors: *RK2*ParD (green), *Cc*PArD (purple), *Mo*ParD3 (yellow) and *Sa*CopG (cyan).

Like other members of the MetJ/CopG/Arc family including its homologs from plasmid RK2, *M. opportunistum* and *C. crescentus*, the *Vc*ParD2 monomers associate into a conserved dimer (Figure 2a,b). Dimer formation buries about 1240 Å^2^ of mainly hydrophobic surface and creates the hydrophobic core of the protein. According to the P(Δ^i^G) value of 0.37 provided by the PISA server (Krissinel & Henrick, 2007), the contact surfaces are somewhat more hydrophobic that the average surface of a soluble protein, in agreement with a dimer that would be stable in solution. Hydrophobic contacts involve contributions from most non-polar side chains: Ile 7 and Leu9 at the end of the N-terminal β-strand, Phe13 and Phe16, in helix α1 and Ile17, Ala29, Val32, Ile33, Ala36, Leu37, Leu39 and Leu40 in helix α2. Most notable is the close contact between the α213 helices of both chains, where Ala36 seems to be essential, any other residue except for glycine at this position resulting in steric crowding. Further stabilization comes via hydrogen bonds, most importantly via pairing of the N-terminal β-strands.

### 3.3. Higher order structure in the crystal

In the crystal, the *Vc*ParD2_N_ dimers are found in a circular arrangement. This results in a torus consisting of eight *Vc*ParD2_2_ dimers (Figure 3). The interface formed by adjacent *Vc*ParD2_2_ dimers buries roughly 600 Å^2^. This interface is highly hydrophilic (PISA P(Δ^i^G) value of 0.91) and is dominated by hydrogen bonds and salt bridges, and as such deviates from typical stable oligomerization interfaces (Figure 4a). The interface is formed by seven amino acid side chains: Arg25, Tyr26, Glu31, Arg34, Arg38, Glu41 and Asn42. These side chains are involved mostly in inter-side chain hydrogen bonds or salt bridges (between Glu41 and Arg25, and between Glu31 and Arg34 as well as Arg38) and cation-π stacking between Arg38 and Tyr26. The only hydrogen bond involving a main chain atom is between the main chain carbonyl of Arg25 and the side chain of Arg34. Of interest is also the close contact between the side chains of Arg38 of both chains, their NH2 atoms being only 3.1 Å apart. This together with the dominance of side chain-side chain interactions (which are entropically less favorable than main chain-main chain interactions) would normally let us conclude that this interface only corresponds to a packing contact and is unlikely to exist in solution.

**Figure 3.**
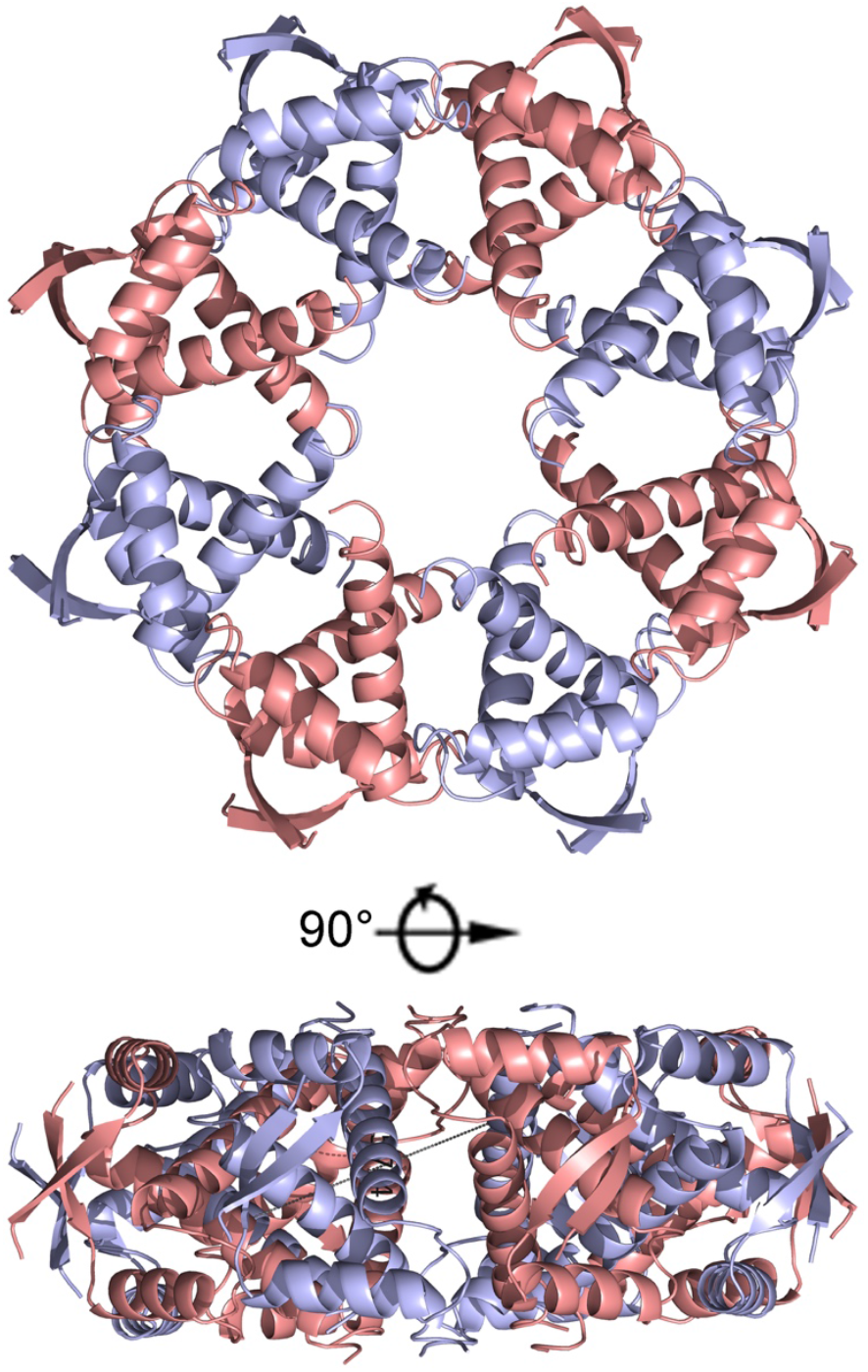
Higher order oligomeric structure of VcParD2_N_. Two orientations of a cartoon representation of the hexadecamer of *Vc*ParD2 as seen in the crystal. *Vc*ParD2 dimers are alternating colored pink and blue.

**Figure 4.**
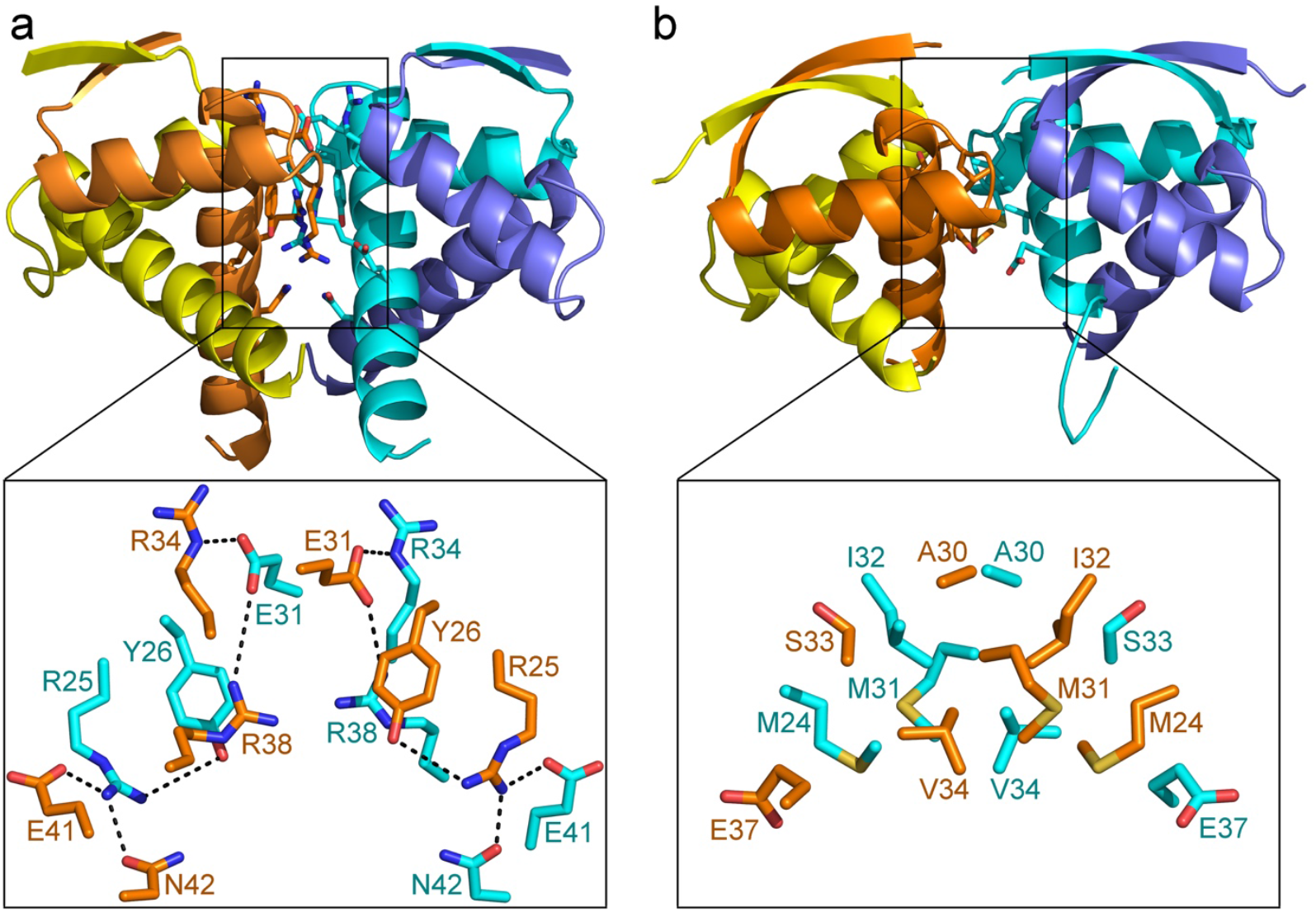
Inter-dimer interface. **a**. Cartoon representation of the *Vc*ParD2 tetramer. Each chain is colored differently. Side chains involved in inter-dimer contacts are shown in stick representation and are shown enlarged at the bottom. The contacts are dominated by hydrogen bonds and salt bridges. **b**. Cartoon representation of the *Sa*CopG tetramer as found in the *Sa*CopG-DNA complex and shown in a similar orientation as *Vc*ParD2. The details of the interacting side chains are again shown enlarged in the bottom of the panel. The inter-dimer contacts involve almost exclusively hydrophobic van de Waals interactions.

The amino acids contributing to this interface are retained in *Cc*ParD, which was previously shown to be a dimer in solution (Dalton & Crosson, 2010). The only substitution among the residues involved in the inter-dimer interface is Asn42, which is a Glu43 in *Cc*ParD. This substitution forces Glu42 of *Cc*ParD into a different conformation than its *Vc*ParD2 equivalent Glu41 in order to separate the two negative charges. This substitution would also result in an interface with a net negative charge of -2. Another potentially relevant difference between *Vc*ParD2 and *Cc*ParD is the reorientation of Arg38 (Arg39 in *Cc*ParD), that now interacts with Glu43 of *Cc*ParD. This further results in an “inwards” movement of Arg26 of *Cc*ParD (Arg25 of *Vc*ParD2) to fill up the space vacated by Arg39. The resulting theoretical interface would then be destabilized by close contacts between the positively charged side chains of Arg26 and Arg39 of the adjacent polypeptide chains.

The hole at the center of the torus has a diameter of about 16 Å. The C-termini of the 16 *Vc*ParD2 N-terminal domains point outwards from the middle of the side surface of the torus. If this arrangement would be composed of intact *Vc*ParD2 chains, the C-terminal IDP regions (residues Leu52-Arg81) would be forced to adopt a fan-shaped ensemble that is limited in its conformational variability. In contrast, in *RK*ParD as well as in CcdA the C-terminal IDP ensemble adopts a wide range of conformations, many of them sterically incompatible with the oligomerization of the *Vc*ParD2 N-terminal domains into the torus observed in the crystal (Madl et al., 2006; Oberer et al., 2007). It is therefore safe to state that the IDP region would provide an entropic penalty for *Vc*ParD2 oligomerization similar to what was observed for the influence of the IDP region of Phd on operator binding (Garcia-Pino et al., 2016).

The *Vc*ParD2 16-mer resembles the crystallographic assembly of *Streptococcus agalactiae* CopG, which forms a spiral structure with inter-dimer contact surfaces roughly corresponding to the inter-dimer interfaces seen in the *Vc*ParD2 16-mer (Figure 4b) (Gomis-Rüth et al., 1998). In contrast to *Vc*ParD2, although similar in size, the oligomerization interface of *Sa*CopG is much more hydrophobic (PISA P(Δ^i^G) value of 0.38) and is retained in the complex of tetrameric *Sa*CopG with its operator (Gomis-Rüth et al., 1998; Costa et al., 2001). Yet, in solution in absence of a DNA ligand, *Sa*CopG behaves as a dimer.

### 3.4. DNA binding site

In analogy with *RK*ParD (Roberts et al., 1993), it can be assumed that *Vc*ParD2 is a DNA binding protein that represses the *parDE2* operon. In general, transcription factors of the ribbon-helix-helix family dock with their N-terminal strands into the major groove of their target DNA. In agreement, the N-terminal β-ribbons present a positively charged electrostatic surface on the *Vc*ParD2 dimer that can complement the negative charges of the DNA backbone.

*Streptococcus agalactiae Sa*CopG is the closest structural relative of *Vc*ParD2 for which a structure of a DNA complex is available. *Sa*CopG makes base pair-specific contacts via the side chains of Arg4 and Thr6, while hydrogen bonds with the DNA backbone are provided by the side chains of Thr8, Lys28 and Ser29 (Gomis-Rüth et al., 1998 - PDB entry 1B01). In *Vc*ParD2, the corresponding residues are Asn4, Ser6, Thr8, Ser29 and Ser31, indicating potential differences in DNA specificity between the two proteins (Supplementary Figure S2a). Unfortunately, the target sequence of *Vc*ParD2 remains currently unknown.

*Sa*CopG binds its operator in a chain-like fashion. Several dimers position themselves next to each other in a way similar as seen in the crystal packing of the free *Sa*CopG structure, with the DNA wrapping around this protein helix (Costa et al., 2001). In contrast to *Sa*CopG, *Vc*ParD2 does not associate into a helical structure, but forms a closed circle. The dimer-dimer association of *Vc*ParD2 is nevertheless very similar to the one of *Sa*CopG bound to DNA (Figure 5a), suggesting that also here crystal packing may reflect associations occurring on the DNA. Indeed, on the toroidal surface of the *Vc*ParD2 crystallographic 16-mer, the N-terminal β-strands stick out as a set of eight parallel and equally spaced ridges that lead to a cogwheel-like arrangement (Figure 3, 5). Figure 5b shows the positioning of an 18 bp fragment with 2 nucleotides overhang taken from a *Sa*CopG-DNA complex and covering two *Sa*CopG dimers on the surface of the *Vc*ParD2 oligomer (Figure 5a). When superimposed upon *Vc*ParD2, the helical axis of the DNA is tangential to the cogwheel surface, and the subsequent β-strand ridges are spaced as such to make it possible for a double-stranded DNA to wrap around the cogwheel. Thus likely, the oligomerization mode of *Vc*ParD2 may reflect how it interacts with its operator.

**Figure 5.**
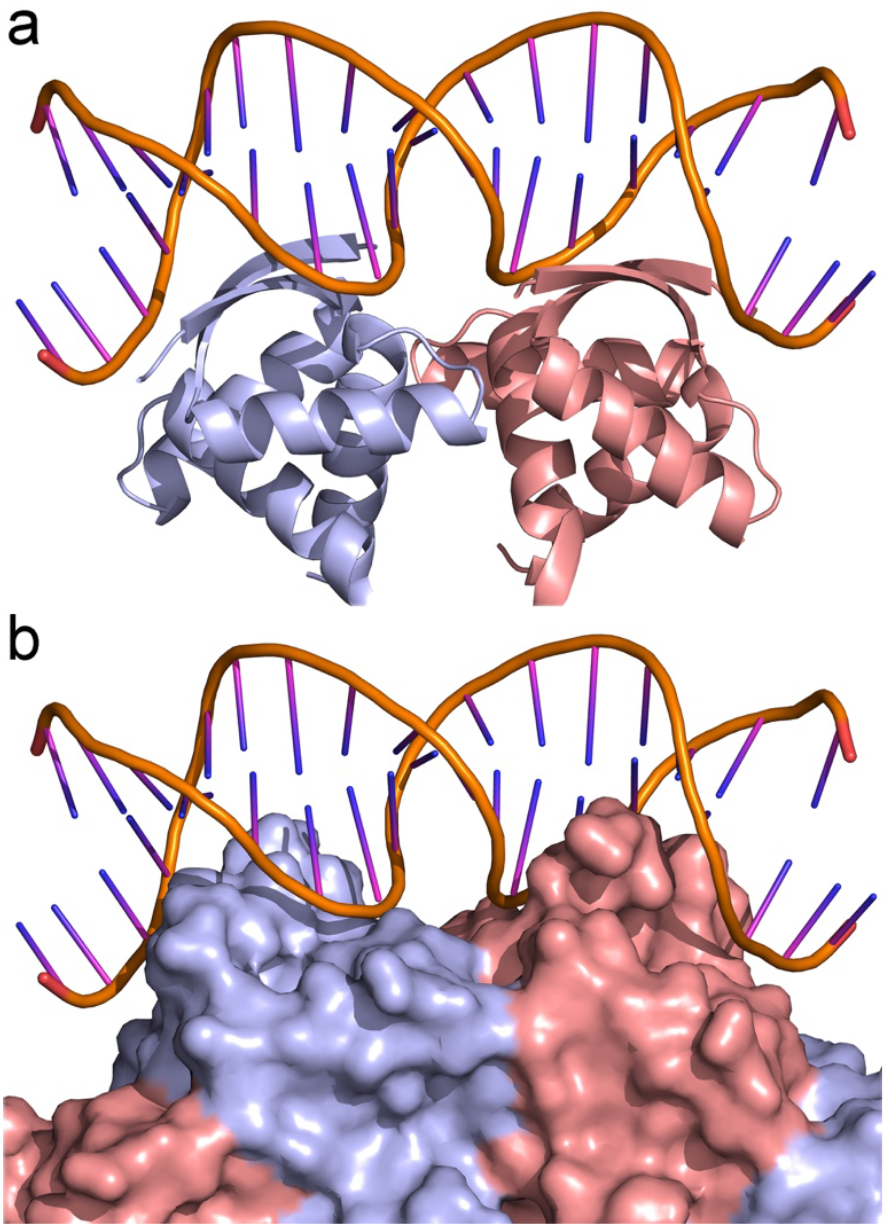
DNA binding site. **a**. Complex of *S. agalactiae Sa*CopG with an 18 bp DNA fragment. **b**. Model of *Vc*ParD2 (surface representation as in Figure 3A) in complex with DNA. The *Sa*CopG tetramer was superimposed in two adjacent dimers of *Vc*ParD2. The DNA fragment bound to *Sa*CopG fits on the surface of *Vc*ParD2 with the N-terminal β-strands inserted into the DNA major groove.

### 3.5. Oligomerization in solution

Despite that there are several arguments that disfavor the existence of the crystalline oligomeric assembly in solution, we did observe already during the purification of *Vc*ParD2 that it elutes on a Superdex Increase 200 10:300 SEC column at an appreciably higher molecular weight than is expected for a simple dimer, even if we take into account that the IDP region would increase its hydrodynamic radius substantially. To evaluate the relevance of the oligomer seen in the crystal, we determined the oligomeric state of *Vc*ParD2 in solution using SEC-MALS and native mass spectrometry.

To get a better estimate of the true oligomeric state in solution and how it may be affected by concentration, we turned to SEC-MALS. A concentration series of *Vc*ParD2 ranging from 18 mg ml^-1^ to 0.1 mg ml^-1^ was injected into a Shodex KW402.5-4F column. For all concentrations, a single peak was observed at an elution volume indicating a higher order oligomer (Figure 6, supplementary Table I, Supplementary Figure S3). The molecular weight determined for the protein eluting in this peak ranges from 100.0.0 ± 0.6 kDa for the highest concentration used to 78.5 ± 0.6 kDa for the lowest concentration used. No additional peaks were observed that can be attributed to a monomer or dimer. Given that the theoretical molecular weight for a *Vc*ParD2 dimer equals 17912 Da and that it can be assumed that the oligomeric state of *Vc*ParD2 is a multiple of dimers, the SEC-MALS data indicate the presence of mainly decamers, likely in equilibrium with dodecamers and octamers.

**Figure 6.**
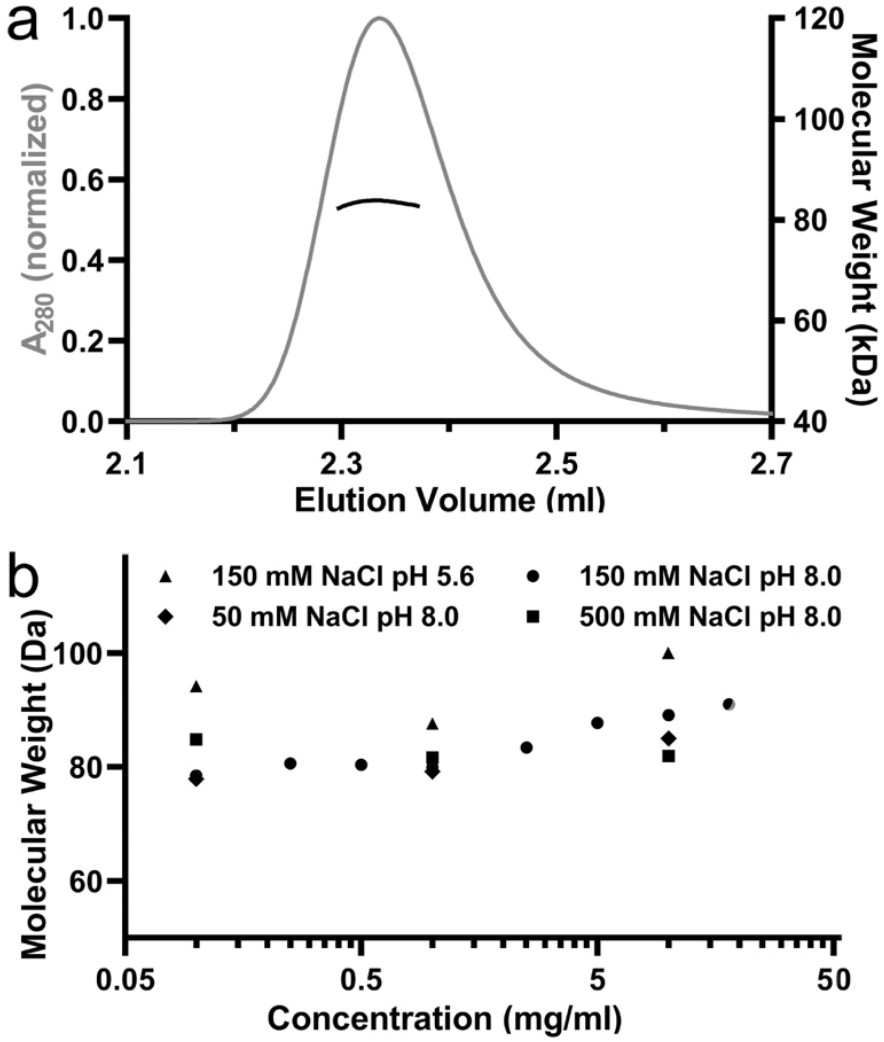
SEC-MALS. **a**. Typical SEC-MALS run (10 mg/ml *Vc*ParD2, 20 mM Tris pH 8.0, 150 mM NaCl, 1 mM TCEP). The observed elution peak and plateau for molecular mass is representative for all conditions tested. **b**. MALS-derived molecular weights as function of concentration and experimental conditions. No clear concentration or ionic strength dependence is observed. Molecular mass values at pH 5.6 are nevertheless systematically higher than for the measurements at pH 8.0.

Little concentration or ionic strength dependence is observed, although lower concentrations tend towards somewhat lower average molecular weights. Similarly, lower pH leads to a somewhat higher average molecular weight. Most important, however, is that even at the highest concentration used, the experimental molecular weight is significantly less than that of a hexadecamer (as suggested from the crystal structure). This deviation from the expected molecular weight is not due to degradation of the C-termini as nESI-TOF mass spectra of the protein sample after performing the SEC-MALS experiments showed the protein to be intact with no signs of degradation.

To further understand the true nature of the higher order complexes present in solution, we performed native mass spectrometry (Figure 7, Supplementary Figure S4). These data nicely mirror the results obtained using SEC-MALS and show that *Vc*ParD2 is predominantly present in solution as a mixture of decamers and dodecamers, and tetramers as a third species. Similar to what is seen in SEC-MALS, lower pH favors the dodecamer assembly. At the lowest concentrations (0.1 mg ml^-1^), some monomers, tetramers and a very small amount of dimers can nevertheless be seen as well, but decamers and dodecamers still dominate, indicating that these are indeed stable species. The stability of the decamers and dodecamers, particularly at high ionic strength, is surprising given the electrostatic nature of the inter-dimer contact interface, as is also the lack of substantial amounts of tetramers, hexamers or octamers. We do not observe the presence of tetradecamers nor the hexadecamer that would be expected from the crystal structure.

**Figure 7.**
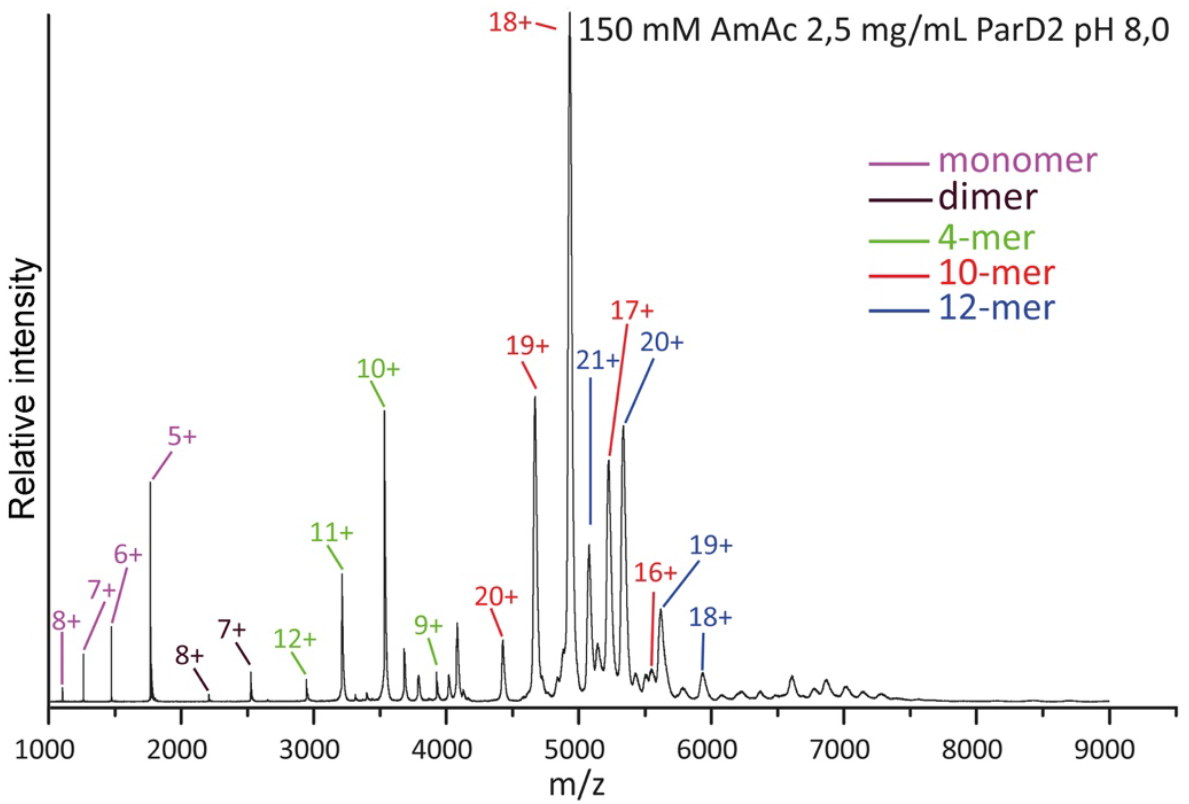
Native mass spectrometry. The native mass spectrum of *Vc*ParD2 (2.5 mg/ml) in 150 mM ammonium acetate pH 8.0 is shown with the major peaks labeled according to their charge state. The major species present are decamers and dodecamers, with smaller amounts of monomers, dimers and tetramers also observed.

### 3.6. Solution SAXS model for the *Vc*ParD2 oligomer

In order to obtain a more detailed picture of the oligomeric species in solution, we turned to small angle X-ray scatter (SAXS). The protein used in this experiment was again confirmed to be intact using nESI-TOF mass spectrometry and is therefore expected to contain a significant fraction of its polypeptide (27%) to be in a disordered ensemble. SAXS data collected at the SWING beamline (Soleil, Saint-Aubin, France) are of high quality up to a *q* value of 0.5 (Figure 8a and Table III). The Guinier plot shows a linear behavior (R^2^ = 0.97), rendering a radius of gyration of 33.64 ± 0.34 Å. The dimensionless Kratky plot of *Vc*ParD2 does, however, not reveal the disordered nature of the C-terminal regions (Supplementary Figure S5). Its bell-shaped curve with a maximum around (1.73; 1.1) agrees with the expected values for a globular particle. The P(r) function shows a nice bell shape and converges smoothly to zero, with a good fit at low *q* angles, indicative of a polydisperse sample (Franke et al., 2017).

**Figure 8.**
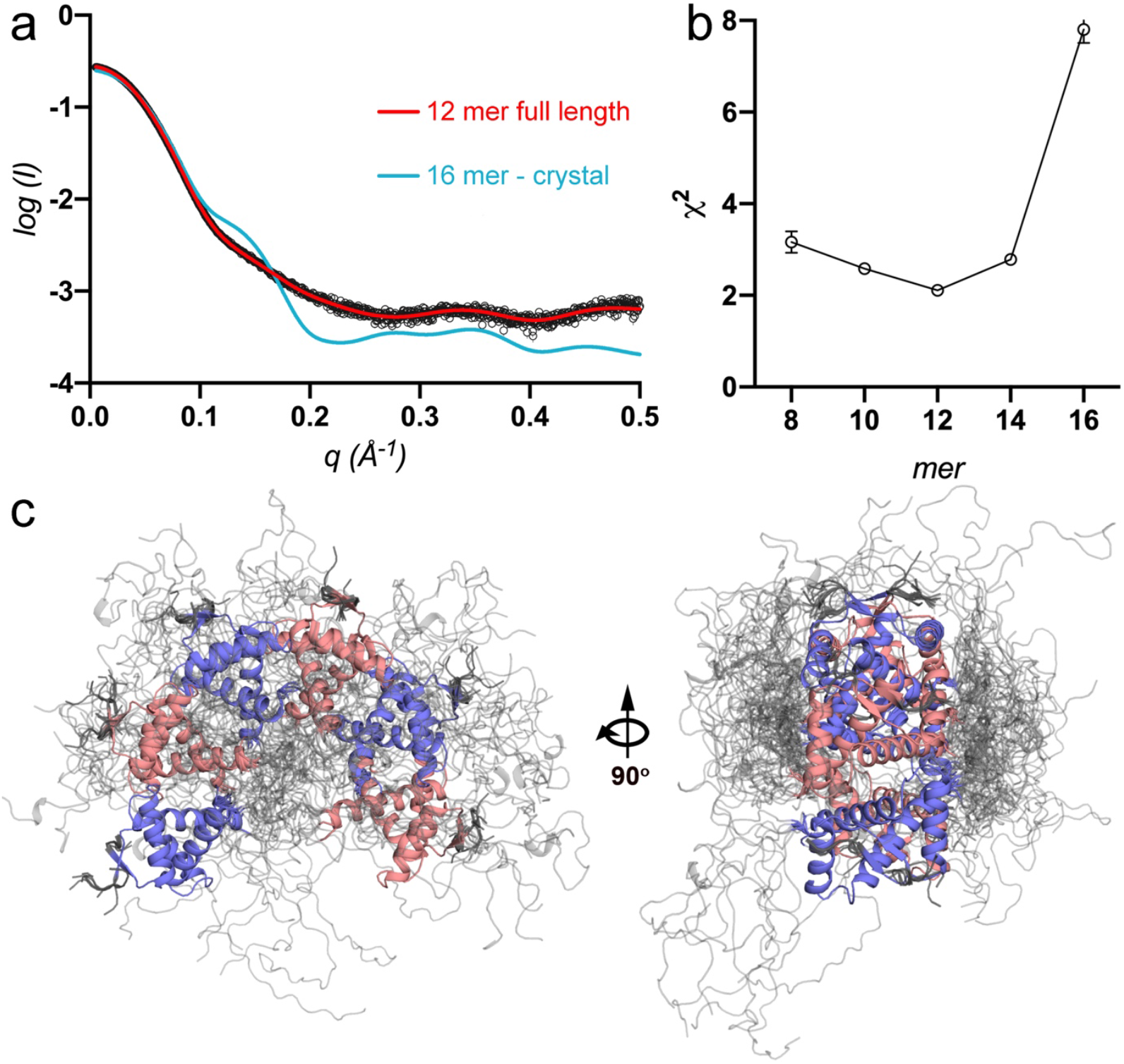
Small angle X-ray scatter. **a**. Experimental SAXS data (black) overlaid with the theoretical scattering curve for a ParD2 12-mer averaged over the 10 best-fitting conformations (red - χ^2^= 2.11) and the curve calculated for the 16-mer crystallographic assembly (cyan). **b**. χ^2^ values for the best fits of 8-, 10-, 12-, 14-mer and 16-mer ensembles. A minimum is seen for a 12-mer ensemble. **c**. Molecular model for the 12-mer ensemble that best represents the scattering data. The folded parts of *Cc*ParD2 dimers are alternately shown in blue and pink. The IDP tails are shown in grey.

**Table 1.**
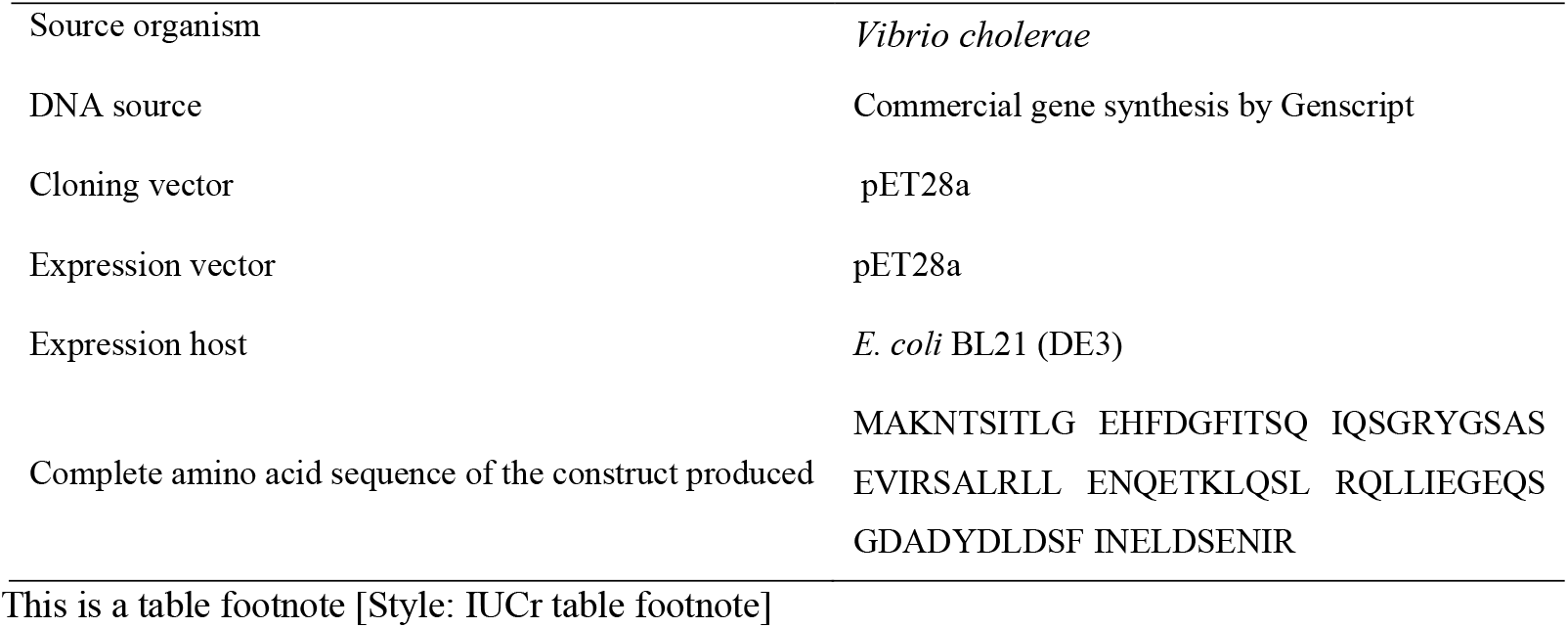
Macromolecule production information. In the primers, indicate any restriction sites, cleavage sites or introduction of additional residues, *e*.*g*. His6-tag, as well as modifications, *e*.*g*. Se-Met instead of Met. [Style: IUCr table headnote]

**Table 2.** Crystallization.

|  |  |
| --- | --- |
| Method | sitting drop |
| Plate type | 96-well IntelliPlate (Hampton Research) |
| Temperature (K) | 293 |
| Protein concentration | 20 mg ml<sup>-1</sup> |
| Buffer composition of protein solution | 20 mM Tris pH 8.0, 150 mM NaC |
| Composition of reservoir solution | 0.2 M Lithium sulfate, 0.1 M MES pH 6.0 and 20 % w/v PEG<br>4000 |
| Volume and ratio of drop | 0.2 µl, 1:1 |
| Volume of reservoir | 100 µl |

**Table 3.**
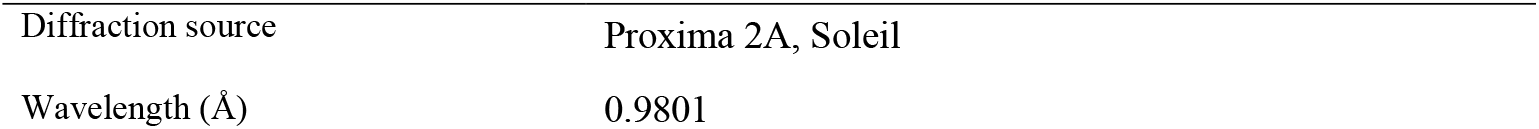
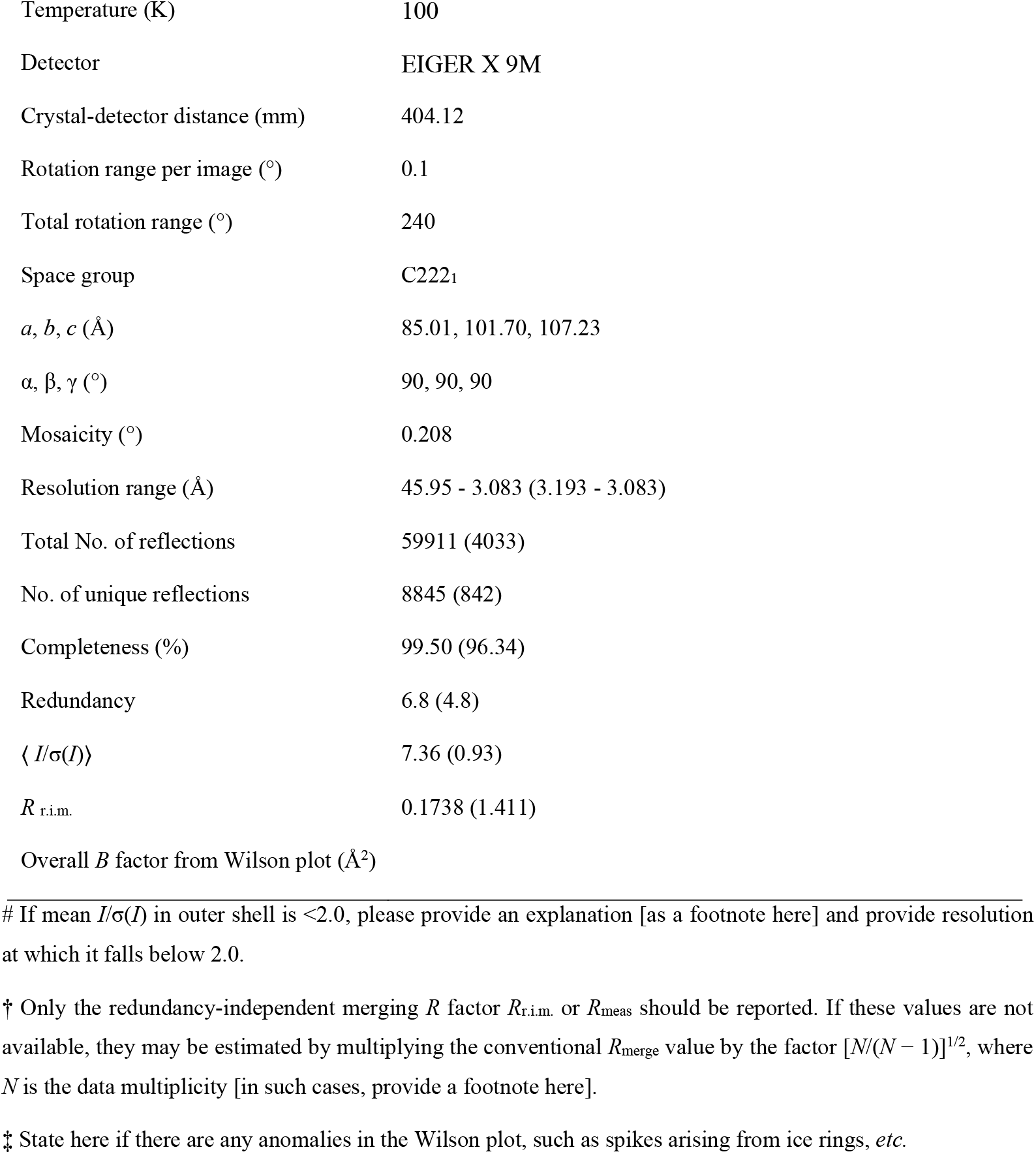
Data collection and processing. Values for the outer shell are given in parentheses.

|  |  |
| --- | --- |
| Diffraction source | Proxima 2A, Soleil |
| Wavelength (Å) | 0.9801 |
| Temperature (K) | 100 |
| Detector | EIGER X 9M |
| Crystal-detector distance (mm) | 404.12 |
| Rotation range per image (°) | 0.1 |
| Total rotation range (°) | 240 |
| Space group | C222<sub>1</sub> |
| *a*, *b*, *c* (Å) | 85.01, 101.70, 107.23 |
| $\alpha$, $\beta$, $\gamma$ (°) | 90, 90, 90 |
| Mosaicity (°) | 0.208 |
| Resolution range (Å) | 45.95 - 3.083 (3.193 - 3.083) |
| Total No. of reflections | 59911 (4033) |
| No. of unique reflections | 8845 (842) |
| Completeness (%) | 99.50 (96.34) |
| Redundancy | 6.8 (4.8) |
| $\langle I/\sigma(I) \rangle$ | 7.36 (0.93) |
| $R_{\text{r.i.m.}}$ | 0.1738 (1.411) |
| Overall *B* factor from Wilson plot (Å<sup>2</sup>) |  |

SAXS data for *Vc*ParD2 antitoxin indicate a molecular mass of approximately 109 kDa from the Bayesian estimate implemented in the ATSAS suite (Franke et al., 2017). The oligomeric state suggested by this molecular mass thus corresponds to a dodecamer. The sample used for these experiments was verified afterwards with nESI-TOF mass spectrometry and was shown to consist only of intact chains. In agreement with this observation, the theoretical SAXS curve calculated for the 16-mer crystallographic assembly does not fit the experimental scattering curve (χ^2^= 28.16). Equally the full-length protein in this hexadecameric assembly with an ensemble where residues 5-47 are fixed and residues 1-4 and 48-81 are given full torsional degree of freedom, still provides a poor fit (χ^2^= 7.80). In this ensemble, the IDP tails tend to plug the central hole, rendering the whole particle more globular. While these solutions are physically possible (there are very few, if any VdW clashes; torsion angles are good; local geometry is perfect), they do not make sense biochemically. Entropic considerations make such a structure, with tails crammed up in the middle, highly unlikely.

We then considered a number of ensembles consisting of full length *Vc*ParD2 chains in 8-, 10-, 12- and 14-mer arrangements. As seen in Figure 8b, the best agreement is obtained for a 12-mer that forms an open fragment of a torus with the relative orientations contacts between adjacent dimers identical as seen in the 16-mer arrangement in the crystal (χ^2^= 2.11). This ensemble features a realistic spread of tails, some of them visiting the central inter-domain region (former central hole), others flopping about on the periphery (Figure 8c). Similar 10-mer and 14-mer models nevertheless fit also reasonably well (χ^2^= 2.58 and 2.78 resp.) and can therefore not be excluded. Hybrid models containing increasing fractions of 10-mer structures added to the 12-mer ensemble do not further improve the fit. In other words, the 12-mer ensemble on its own is sufficient to explain the scattering data and is most likely the dominant species present in solution.

## 4. Discussion

The folded domains of *Vc*ParD2 form a partial doughnut structure in solution that is stabilized via strong electrostatic interactions. While not recognized as such by the algorithm of the PISA server (Krissinel & Henrick, 2007), this association is stable and maintained even at low protein concentrations. For over four decades, hydrophobicity next to complementarity is believed to be the major factor stabilizing protein-protein association (Chothia & Janin, 1975). Contact surfaces in protein oligomers differ from the rest of the subunit surface in that they are enriched in hydrophobic side-chains and have a low density of inter-subunit hydrogen bonds (Janin et al., 1988; Lo Conte et al., 1999). For small dimeric proteins, the dimer interface often corresponds to the hydrophobic core and dimerization is essential to form a stable structure.

In contrast, high affinity electrostatic interactions are known between IDPs, e.g. the disordered complex formed between histone H1 and its chaperone prothymosin-α (Borgia et al., 2018). Similar poly-electrolytic interactions are also postulated to drive the formation of membrane-less organelles via liquid-liquid phase transitions (Brangwynne et al., 2015; Schuler et al., 2020). On the other hand, to find a stable association of globular proteins dominated by side chain-side chain hydrogen bonds and electrostatic interactions is highly unusual. To our knowledge, no other examples of such complexes are known, the *Vc*ParD2 oligomer thus representing a unique case. The stability of this oligomer is even more surprising when one considers that the amino acids involved in this interaction are highly conserved in *Cc*ParD from *C. crescentus*, the only difference being the substitution of an Asn for Glu in *Cc*ParD. The latter would bury two negative charges at the inter-dimer interface, creating highly unstable situation.

In the crystal, the globular domain of *Vc*ParD2 forms a closed doughnut-shaped structure in absence of its IDP tails, which most likely degraded in the two months that were required for crystals to appear. In absence of any cooperativity, one would expect that with identical interactions between all dimers, in solution also a complete circle would be formed. We could demonstrate nevertheless that in solution, the doughnut is incomplete and that the assembly consists of 5-6 *Vc*ParD2 dimers instead of 8. This implies the existence of negative cooperativity that prevents the formation of the closed structure. Likely this negative cooperativity originates from the IDP tails, which in a full 16-mer assembly would hinder each other’s freedom and create an entropic barrier that prevents an oligomer larger than a 12-mer to form. A similar entropic exclusion principle was previously observed for the binding of Phd to its operator (Garcia-Pino et al., 2016). Two copies of the Phd dimer need to bind at adjacent sites, but this is prevented due to entropic exclusion of the IDP tails. Only when Doc binds and folds these IDP tails, operator binding can proceed with high affinity and the *phd/doc* operon is repressed. This mechanism is further related to the action of entropic bristles, IDP tails that can act as solubilizers to prevent aggregation (Santner et al., 2012), tune prion nucleation (Michiels et al., 2020) and in general tune the energy landscape of proteins with respect to protein assemblies and ligand binding and association (Keul et al., 2018; Niemeyer et al., 2020).

The similarities in higher-order association between *Vc*ParD2 and *Sa*CopG from *Streptococcus agalactiae* suggest a mechanism for DNA binding. *Vc*ParD2 and *Sa*CopG make higher order contacts via the same spatial surface, although this surface for *Sa*CopG is substantially more hydrophobic. This association in *Sa*CopG leads to an extended DNA binding surface where multiple *Sa*CopG dimers dock next to each other on the operator. In many TA systems, the antitoxin often binds to two or more binding sites on the operator. In most cases, the toxin increases antitoxin-DNA affinity by bridging adjacent antitoxin dimers (Garcia-Pino et al., 2010; Vandervelde et al., 2016; Xue et al., 2020), or by otherwise stabilizing the DNA-binding assembly the antitoxin (Bøggild et al., 2012; Qian et al., 2019; Jurėnas et al., 2019). Higher toxin to antitoxin ratios lead to de-repression via the formation of toxin-antitoxin complexes with altered stoichiometry. For antitoxins that bind only to a single site, regulation is less complex and the toxin only serves to weaken operator binding (Brown et al., 2013; Turnbull & Gerdes, 2018; Winter et al., 2018; Manav et al., 2019). For *parDE* modules, the mechanism of transcription regulation is not known. Early studies on the *parDE* system present on plasmid RK2 indicated that the antitoxin is sufficient to repress the operon and likely binds on two adjacent palindromes (Roberts et al., 1993). Although the operator of the *VcparDE2* operon is not known, the higher order association of *Vc*ParD2 dimers suggest cooperative binding to the DNA in a way similar to what is observed for *Sa*CopG. To what extent the corresponding toxin influences this interaction remains currently unknown.

**Table 4.** Structure solution and refinement.

|  |  |
| --- | --- |
| Resolution range (Å) | 45.95 - 3.08 (3.19 - 3.08) |
| Completeness (%) | 99.50 (96.34) |
| $\sigma$ cutoff | none |
| No. of reflections, working set | 8845 (842) |
| No. of reflections, test set | 442 (42) |
| Final $R_{\text{cryst}}$ | 0.27 (0.35) |
| Final $R_{\text{free}}$ | 0.29 (0.36) |
| No. of non-H atoms | 2856 |
| Protein | 2856 |
| Total | 2856 |
| R.m.s. deviations |  |
| Bonds (Å) | 0.001 |
| Angles (°) | 0.30 |
| Average $B$ factors (Å<sup>2</sup>) | 84.57 |
| Protein | 84.57 |
| Ramachandran plot |  |
| Most favoured (%) | 95.74 |
| Allowed (%) | 3.99 |
| Disallowed (%) | 0.27 |

Statistics for the highest-resolution shell are shown in parentheses.

**Table 5.**
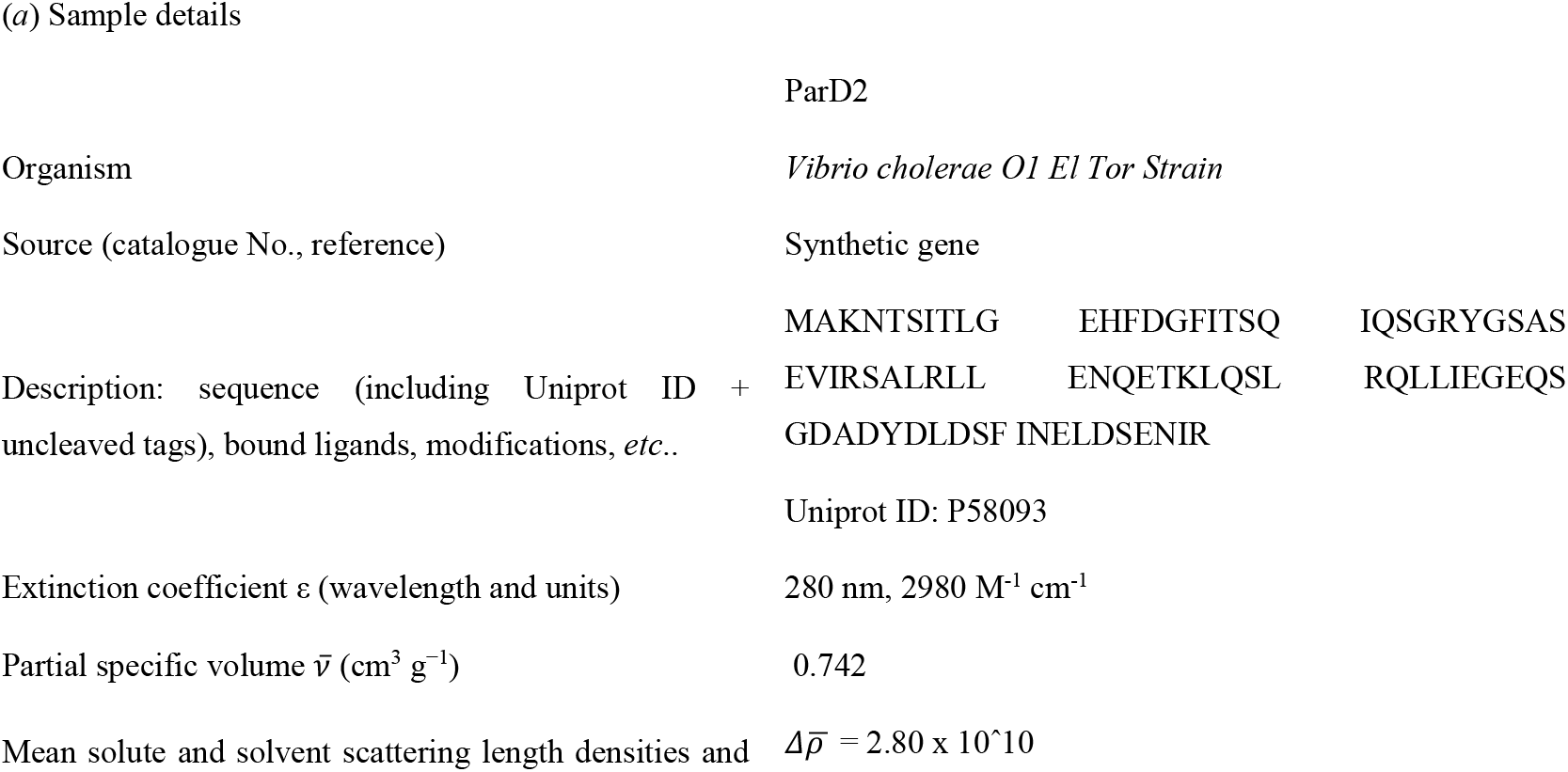
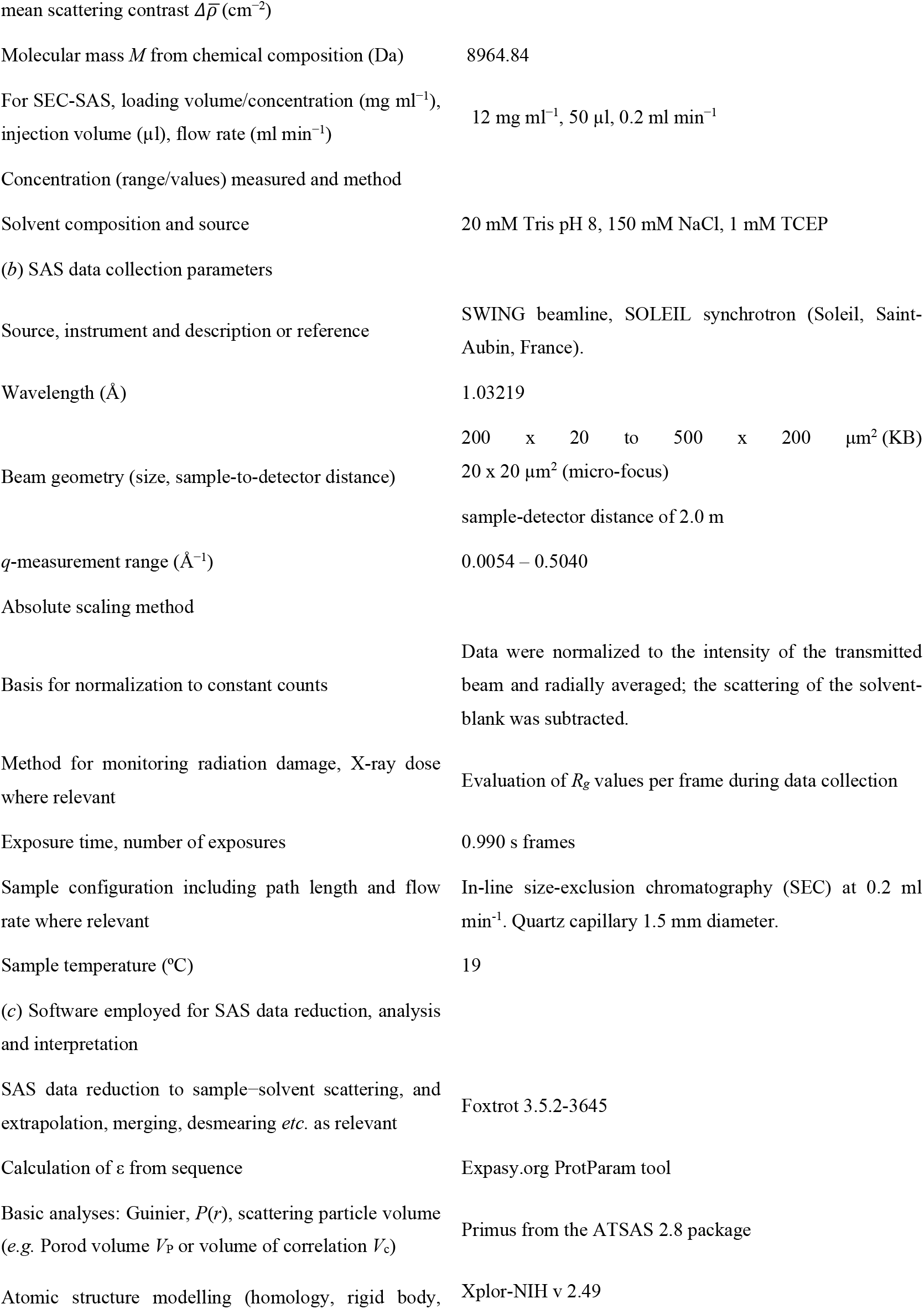
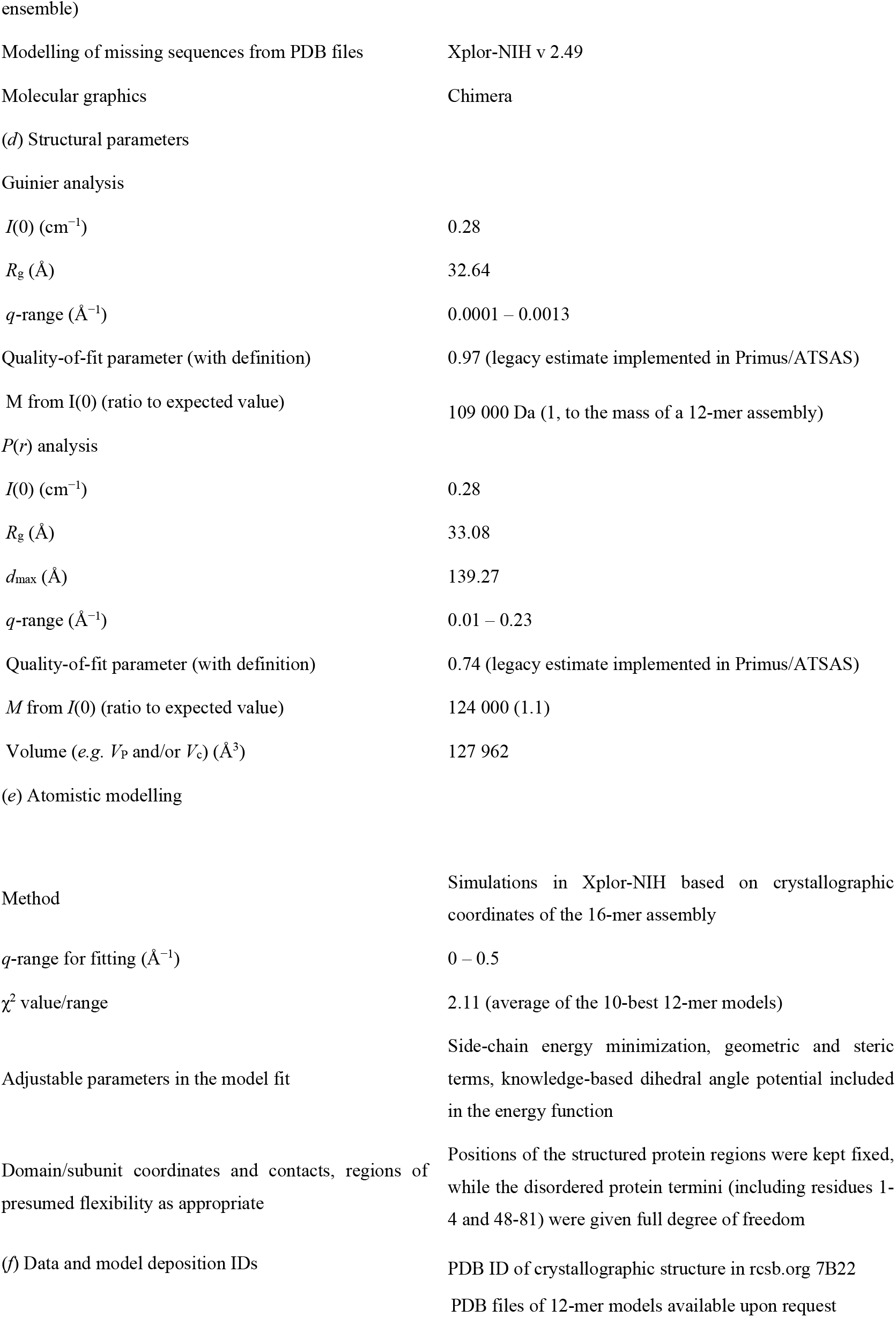
SAS experimental details. This table has been prepared according to guidelines published by Trewhella *et al*. (2017), *Acta Cryst*. D**73**, 710-728, https://doi.org/10.1107/S2059798317011597.

**Table 6.** Structural homologs of the *Vc*ParD2 N-terminal domain picked up in a Dali search. This is a table headnote (style: IUCr table headnote)

| Protein | Organism | PDB entry | Dali Z-score | rmsd (Å) | Seq. ID<br>(N-terminal domain) | Number of common<br>Cα atoms | Reference |
| --- | --- | --- | --- | --- | --- | --- | --- |
| *CcParD* | *Caulobacter crescentus* | 3kxe | 7.5 | 0.7 | 63% | 47 | Dalton & Crosson, 2010 |
| *SoCopA* | *Shewanella oneidensis* | 6iya | 6.2 | 1.8 | 20% | 46 | Zhao et al., 2019 |
| *MoParD3* | *Mesorhizobium opportunistum*<br>*WSM2075* | 5ceg | 6.1 | 1.7 | 17% | 43 | Aakre et al., 2015 |
| *SaCopG* | *Streptococcus agalactiae* | 2cpg | 5.6 | 1.8 | 23% | 43 | Gomis-Rüth et al., 1998 |
| *Arc* | *Salmonella* bacteriophage P22 | 1arq | 5.4 | 1.9 | 7% |  | Bonvin et al., 1994 |
| *EcAtaR* | *Escherichia coli* | 6ajn<br>6gto | 5.2 | 2.1 | 9% | 44 | Yashiro et al., 2019;<br>Jurėnas et al., 2019 |
| *RKParD* | *E. coli* RK2 plasmid | 2an7 | 4.4 | 2.1 | 8% | 33 | Oberer et al., 2007 |

This is a table footnote (style: IUCr table footnote)

## Acknowledgements

The authors thank Sarah Haesaerts for technical assistance, Didier Vertommen (UCL) for nESI-TOF mass spectrometry and the beamline staff of the Soleil macromolecular crystallography and SAXS beamlines for help during data collection and processing.

## References

Aakre C.D., Herrou J., Phung T.N., Perchuk B.S., Crosson S., Laub M.T. (2015) Evolving new protein-protein interaction specificity through promiscuous intermediates. Cell 16, 594–606.

Afonine P.V., Grosse-Kunstleve R.W., Echols N., Headd J.J., Moriarty N.W., Mustyakimov M., Terwilliger T.C., Urzhumtsev A., Zwart P.H., Adams P.D. (2012) Towards automated crystallographic structure refinement with phenix.refine. Acta Crystallogr. D68, 352–367.

Bøggild A., Sofos N., Andersen K.R., Feddersen A., Easter A.D., Passmore L.A., Brodersen D.E. (2012) The crystal structure of the intact E. coli RelBE toxin-antitoxin complex provides the structural basis for conditional cooperativity. Structure 20, 1641–1648.

Bonvin A.M., Vis H., Breg J.N., Burgering M.J., Boelens R., Kaptein R. (1994) Nuclear magnetic resonance solution structure of the Arc repressor using relaxation matrix calculations. J. Mol. Biol. 236, 328–341.

Borgia A., Borgia M.B., Bugge K., Kissling V..M, Heidarsson P.O., Fernandes C.B., Sottini A., Soranno A., Buholzer K.J., Nettels D., Kragelund B.B., Best R.B., Schuler B. (2018) Extreme disorder in an ultrahigh-affinity protein complex. Nature 555, 61–66.

Brangwynne C.P., Tompa P., Pappu R.V. (2015) Polymer physics of intracellular phase transitions. Nat. Physics 11, 899–904.

Brown B.L., Lord D.M., Grigoriu S., Peti W., Page R. (2013) The Escherichia coli toxin MqsR destabilizes the transcriptional repression complex formed between the antitoxin MqsA and the mqsRA operon promoter. J. Biol. Chem. 288, 1286–1294.

Chothia C., Janin J. (1975) Principles of protein-protein recognition. Nature 256, 705–708.

contents of proteins, DNA, and protein-nucleic acid complex crystals. Protein Sci. 12, 1865–1871.

Costa M., Solà M., del Solar G., Eritja R., Hernández-Arriaga A.M., Espinosa M.E., Gomis-Rüth F.X., Coll M. (2001) Plasmid transcriptional repressor CopG oligomerises to render helical superstructures unbound and in complexes with oligonucleotides. J. Mol. Biol. 310, 403–417.

Dalton K.M., Crosson S. (2010) A conserved mode of protein recognition and binding in a ParD-ParE toxin-antitoxin complex. Biochemistry 49, 2205–2215.

De Jonge N., Buts L., Garcio-Pino, A., Buts, L., Haesaerts S., Zangger K., De Greve H., Loris R. (2009) Rejuvenation of CcdB-poisoned gyrase by an intrinsically disordered protein domain. Mol. Cell 35, 154–163.

Díaz-Orejas R., Espinosa M., Yeo C.C. (2017) The importance of the expendable: toxin–antitoxin genes in plasmids and chromosomes. Front. Microbiol. 8, 1479.

Emsley P., Lohkamp B., Scott W.G., Cowtan K. (2010) Features and development of Coot. Acta Crystallogr. D66, 486–501.

Fineran P.C. (2019) Resistance is not futile: bacterial ‘innate’ and CRISPR-Cas ‘adaptive’ immune systems. Microbiology 165, 834–841.

Franke D., Petoukhov M.V., Konarev P.V., Panjkovich A., Tuukkanen A., Mertens H.D.T., Kikhney A.G., Hajizadeh N.R., Franklin J.M., Jeffries C.M., Svergun D.I. (2017) ATSAS 2.8: a comprehensive data analysis suite for small-angle scattering from macromolecular solutions. J. Appl. Crystallogr. 50, 1212–1225.

Garcia-Pino A., Balasubramanian S., Wyns L., Gazit E., De Greve H., Magnuson R.D., Charlier D., van Nuland N.A.J., Loris R. (2010) Allostery and intrinsic disorder mediate transcription regulation by conditional cooperativity. Cell 142, 101–111.

Garcia-Pino A., De Gieter S., Talavera A., De Greve H., Efremov R.G., Loris R. (2016) An intrinsically disordered entropic switch determines allostery in Phd/Doc regulation. Nat. Chem. Biol. 12, 490–496.

Gasteiger E., Hoogland C., Gattiker A., Duvaud S., Wilkins M.R., Appel R.D., Bairoch A. (2005) Protein identification and analysis tools on the ExPASy Server. In The Proteomics Protocols Handbook, John M. Walker (ed), Humana Press. pp. 571–607

Gerdes K., Rasmussen P.B., Molin S. (1986) Unique type of plasmid maintenance function: postsegregational killing of plasmid-free cells. Proc. Natl. Acad. Sci. USA 83, 3116–3120.

Gomis-Ruth F.X., Sola M., Acebo P., Parraga A., Guasch A., Eritja R., Gonzalez A., Espinosa M., del Solar G., Coll M. (1998) The structure of plasmid-encoded transcriptional repressor CopG unliganded and bound to its operator. EMBO J. 17, 7404–7415.

Hadži S., Garcia-Pino A., Haesaerts S., Jurėnas D., Gerdes K., Lah J., Loris R. (2017) Ribosome-dependent Vibrio cholerae mRNAse HigB2 is regulated by a β-strand sliding mechanism. Nucleic Acids Res. 45, 4972–4983.

Holm L. (2020) DALI and the persistence of protein shape. Protein Sci. 29, 128–140.

Hõrak R., Tamman H. (2017) Desperate times call for desperate measures: benefits and costs of toxin-antitoxin systems. Curr. Genet. 63, 69–74.

Janin J.,Miller S., Chothia C. (1988) Surface, subunit interfaces and interior of oligomeric proteins. J. Mol. Biol. 204, 155–164.

Jiang Y., Pogliano J., Helinski D.R., Konieczny I. (2002) ParE toxin encoded by the broad-host-range plasmid RK2 is an inhibitor of Escherichia coli gyrase. Mol. Microbiol. 44, 971–979.

Jurėnas D., Van Melderen L., Garcia-Pino A. (2019) Mechanism of regulation and neutralization of the AtaR-AtaT toxin-antitoxin system. Nat. Chem. Biol. 15, 285–294.

Kantardjieff K.A., Rupp B. (2003) Matthews coefficient probabilities: Improved estimates for unit cell

Kabsch W. (2010) XDS. Acta Crystallogr. D66, 125–132.

Keul N.D., Oruganty K., Schaper Bergman E.T., Beattie N.R., McDonald W.E., Kadirvelraj R. Gross M.L., Phillips R.S., Harvey S.C., Wood, Z. A. (2018). The entropic force generated by intrinsically disordered segments tunes protein function. Nature 563, 584–588.

Krissinel E., Henrick K. (2007) Inference of macromolecular assemblies from crystalline state. J. Mol. Biol. 372, 774–797.

Kumar P., Issac B., Dodson E.J., Turkenburg J.P., Mande S.C. (2008) Crystal structure of Mycobacterium tuberculosis YefM antitoxin reveals that it is not an intrinsically unstructured protein. J. Mol. Biol. 383, 482–493.

Legrand P. (2017) XDSME: XDS Made Easier (2017) GitHub repository, https://github.com/legrandp/xdsme DOI 10.5281/zenodo.837885.

LeRoux M., Culviner P.H., Liu Y.J., Littlehale M.L., Laub M.T. (2020) Stress can induce transcription of toxin-antitoxin systems without activating toxin. Mol. Cell 79, 280-292.e8.

Lo Conte L., Chothia C., Janin J. (1999) The atomic structure of protein-protein recognition sites. J. Mol. Biol. 285, 2177–2198.

Lopatina A., Tal N., Sorek R. (2020) Abortive infection: bacterial suicide as an antiviral immune strategy. Ann. Rev. Virol. 7, 17.1-17.14.

Loris R., Garcia-Pino A. (2014) Disorder and dynamics-based regulatory mechanisms in toxin-antitoxin modules. Chem. Rev. 114, 6933–6947.

Madl T., Van Melderen L., Mine N., Respondek M., Oberer M., Keller W., Khatai L., Zangger K. (2006) Structural basis for nucleic acid and toxin recognition of the bacterial antitoxin CcdA. J. Mol. Biol. 364, 170–185.

Manav M.C., Turnbull K.J., Jurėnas D., Garcia-Pino A., Gerdes K., Brodersen D.E. (2018) The E. coli HicB antitoxin contains a structurally stable helix-turn-helix DNA binding domain. Structure 27, 1675-1685.e3

McCoy A.J. (2007) Solving structures of protein complexes by molecular replacement with Phaser. Acta Crystallogr. D63, 32–41.

Michiels E., Liu S., Gallardo R., Louros N., Mathelié-Guinlet M., Dufrêne Y., Schymkowitz J., Vorberg I., Rousseau F. (2020) Entropic bristles tune the seeding efficiency of prion-nucleating fragments. Cell Rep. 30, 2834–2845.

Niemeyer M., Moreno Castillo E., Ihling C.H., Iacobucci C., Wilde V., Hellmuth A., Hoehenwarter W., Samodelov S.L., Zurbriggen M.D., Kastritis P.L., Sinz A, Calderón Villalobos L.I.A. (2020) Flexibility of intrinsically disordered degrons in AUX/IAA proteins reinforces auxin co-receptor assemblies. Nat. Commun. 11, 2277.

Oberer M., Zangger K., Gruber K., Keller W. (2007) The solution structure of ParD, the antidote of the ParDE toxin antitoxin module, provides the structural basis for DNA and toxin binding. Protein Sci. 16, 1676–1688.

Page R., Peti W. (2016) Toxin-antitoxin systems in bacterial growth arrest and persistence. Nat. Chem. Biol. 12, 208–214.

Pandey D.P., Gerdes K. (2005) Toxin–antitoxin loci are highly abundant in free-living but lost from host-associated prokaryotes. Nucleic acids Res. 33, 966–976.

Porod, G. (1982). Small-Angle X-ray Scattering, edited by O. Glatter & O. Kratky, ch. 2. London: Academic Press.

Qian H., Yu H., Li P., Zhu E., Yao Q., Tai C., Deng Z., Gerdes K., He X., Gan J., Ou H.Y. (2019) Toxin-antitoxin operon kacAT of Klebsiella pneumoniae is regulated by conditional cooperativity via a W-shaped KacA-KacT complex. Nucleic Acids Res. 47, 7690–7702.

Roberts R.C., Helinski D.R. (1992) Definition of a minimal plasmid stabilization system from the broad-host-range plasmid RK2. J. Bacteriol. 174, 8119–8132.

Roberts R.C., Sprangler C., Helinski D.R. (1993) Characteristics and significance of DNA binding activity of plasmid stabilization protein ParD from the broad host-range plasmid RK2. J. Biol. Chem. 268, 27109–27117.

Ronneau S., Helaine S. (2019) Clarifying the link between toxin-antitoxin modules and bacterial persistence. J. Mol. Biol. 431, 3462–3471.

Santner A.A., Croy C.H., Vasanwala F.H., Uversky V.N., Van Y.-Y.J., Dunker A.K. (2102) Sweeping away protein aggregation with entropic bristles: intrinsically disordered protein fusions enhance soluble expression. Biochemistry 51, 7250–7262.

Schuler B., Borgia A., Borgia M.B., Heidarsson P.O., Holmstrom E.D., Nettels D., Sottini A. (2020) Binding without folding - the biomolecular function of disordered polyelectrolyte complexes. Curr. Opin. Struct. Biol. 60, 66–76.

Schwieters C.D., Bermejo G.A.J., Clore, G.M. (2018) Xplor-NIH for molecular structure determination from NMR and other data sources. Prot. Sci. 27, 26–40.

Schwieters C.D., Clore G.M. (2014) Using small angle solution scattering data in Xplor-NIH structure calculations. Prog. Nucl. Magn. Reson. Spectrosc. 80, 1–11.

Shao Y., Harrison E.M., Bi D., Tai C., He X., Ou H.-Y., Rajakumar K., Deng Z. (2011) TADB: a web-based resource for Type 2 toxin-antitoxin loci in bacteria and archaea. Nucleic Acids Res. 39, D606–D611.

Song S., Wood T. (2020a) Toxin/Antitoxin system paradigms: toxins bound to antitoxins are not likely activated by preferential antitoxin degradation. Adv. Biosyst. 4, e1900290.

Song S., Wood T. (2020b) A primary physiological role of toxin/antitoxin systems is phage inhibition. Front. Microbiol. 11, 1895.

Sterckx Y.G., De Gieter S., Zorzini V., Hadži S., Haesaerts S., Loris R., Garcia-Pino A. (2015) An efficient method for the purification of proteins from four distinct toxin-antitoxin modules. Protein Expr. Purif. 108, 30–40.

Sterckx Y.G., Jové T., Shkumatov A.V., Garcia-Pino A., Geerts L., De Kerpel M., Lah J., De Greve H., Van Melderen L., Loris R. (2016) A unique hetero-hexadecameric architecture displayed by the Escherichia coli O157 PaaA2-ParE2 antitoxin-toxin complex. J. Mol. Biol. 428, 1589–1603.

Sterckx Y.G., Volkov A.N., Vranken W.F., Kragelj J., Jensen M.R., Buts L., Garcia-Pino A., Jové T., Van Melderen L., Blackledge M., van Nuland N.A., Loris R. (2014) Small-angle X-ray scattering- and nuclear magnetic resonance-derived conformational ensemble of the highly flexible antitoxin PaaA2. Structure 22, 854–865.

Szekeres S., Dauti M., Wilde C., Mazel D., Rowe-Magnus D.A. (2007) Chromosomal toxin-antitoxin loci can diminish large-scale genome reductions in the absence of selection. Mol. Microbiol. 63, 1588–1605.

Talavera A., Tamman H., Ainelo A., Konijnenberg A., Hadži S., Sobott F., Garcia-Pino A., Hõrak R., Loris R. (2019) A dual role in regulation and toxicity for the intrinsically disordered N-terminus of the toxin GraT. Nat. Commun. 10, 972.

Turnbull K.J., Gerdes K. (2017) HicA toxin of Escherichia coli derepresses hicAB transcription to selectively produce HicB antitoxin. Mol. Microbiol. 104, 781–792.

Vandervelde A., Drobnak I., Hadži S., Sterckx Y.G.J., Welte T., De Greve H., Charlier D., Efremov R., Loris R., Lah J. (2017) Molecular mechanism governing ratio-dependent transcription regulation in the ccdAB operon. Nucleic Acids Res. 45, 2937–2950.

Winter A.J., Williams C., Isupov M.N., Crocker H., Gromova M., Marsh P., Wilkinson O.J., Dillingham M.S., Harmer N.J., Titball R.W., Crump M.P. (2018) The molecular basis of protein toxin HicA-dependent binding of the protein antitoxin HicB to DNA. J. Biol. Chem. 293, 19429–19440.

Xue L., Yue J., Ke J., Khan M.H., Wen W., Sun B., Zhu Z., Niu L. (2020) Distinct oligomeric structures of the YoeB-YefM complex provide insights into the conditional cooperativity of type II toxin-antitoxin system. Nucleic Acids Res. 48, 10527–10541.

Yao J., Guo Y., Wang P., Zeng Z., Li B., Tang K., Liu X., Wang X. (2018) Type II toxin/antitoxin system ParE(SO) /CopA(SO) stabilizes prophage CP4So in Shewanella oneidensis. Environ. Microbiol. 20, 1224–1239.

Yashiro Y., Yamashita S., Tomita K. (2019) Crystal structure of the enterohemorrhagic Escherichia coli AtaT-AtaR toxin-antitoxin complex. Structure 27, 476-484.e3.

Yuan J, Sterckx Y, Mitchenall LA, Maxwell A, Loris R, Waldor MK. (2010) Vibrio cholerae ParE2 poisons DNA gyrase via a mechanism distinct from other gyrase inhibitors. J. Biol. Chem. 285, 40397–40408.

Yuan J., Yamaichi Y., Waldor M.K. (2011) The three Vibrio cholerae chromosome II-encoded ParE toxins degrade chromosome I following loss of chromosome II. J. Bacteriol. 193, 611–619.

Zhao R., Li Q., Zhang J., Li F., Yao J., Zhang J., Liu L., Wang X., Zhang X. (2019) Structure and allosteric coupling of type antitoxin CopA(SO). Biochem. Biophys. Res. Commun. 514, 1122–1127.

