## Supplementary Data for "Entropic pressure controls oligomerization of *Vibrio cholerae* ParD2 antitoxin"

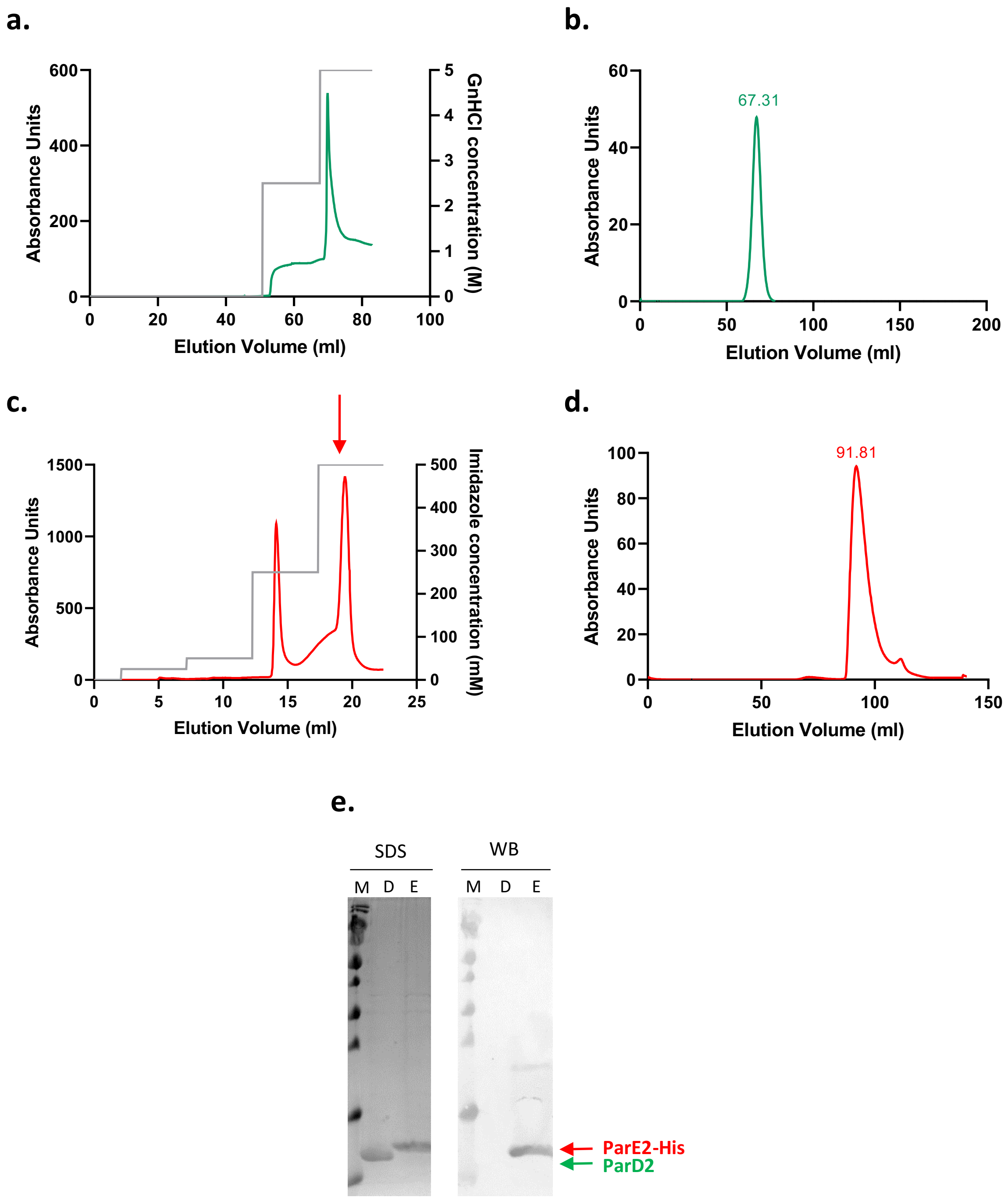


**Supplementary Figure S1.** Purification of *Vc*ParD2 and *Vc*ParE2. **A.** Denaturant-induced dissociation of the *Vc*ParD2:*Vc*ParE2-His complex, resulting in free *Vc*ParD2 and *Vc*ParE2-His trapped on a Ni-NTA column. *Vc*ParD2 from the HisTrap column with a step-gradient of GnHCl. The thus obtained *Vc*ParD2 is refolded by dialysis in 20 mM Tris pH 8, 150 mM NaCl and concentrated to a volume of 2.5 ml **B.** SEC cleaning of the concentrated *Vc*ParD2 on a Superdex 200 16:60 column equilibrated with 20 mM Tris pH 8, 150 mM NaCl. **C.** Elution of on-column refolded *Vc*ParE2-His from the Ni-NTA column using a step-gradient of imidazole. While the first peak still contains some contaminants, the second peak is essentially pure and is concentrated to a volume of 2.5 ml. **D.** SEC cleaning of the concentrated *Vc*ParE2-His on a Superdex 75 16:60 column equilibrated with 20 mM Tris pH 8, 150 mM NaCl, 1 mM TCEP. **E.** SDS-PAGE and corresponding anti-His-tag Western Blot of the pure *Vc*ParD2 and *Vc*ParE2-His proteins after SEC.

**a.**

```
     VcParD2 -------LEEEEELLLLHHHHHHHHHHHLLLLLLHHHHHHHHHHHHHHHHHHLLLLLL---------------------------------------
VcParD2      -----maKNTSITLGEHFDGFITSQIQSGRYGSASEVIRSALRLLENQETKLQSLRqlliegeqsgdadydldsfineldsenir-----------
CcParD  3kxe ----maskNTSVVLGDHFQAFIDSQVADGRYGSASEVIRAGLRLLEENEAKLAALRAALIEGEESGFIEDFDFDAFIEERSRA-------------
MoParD3 5ceg --masnvEKMSVAVTPQQAAVMREAVEAGEYATASEIVREAVRDWLAKRELRHDDIRRLRQLWDEGKASGRPEPVDFDALRKEARQKLTEvppngr
RK2ParD 2an7 ------mSRLTIDMTDQQHQSLKALAAL-QKTI-KQYALERLFPGDADADQAWQELKTMLGNRINDGLAGKVSTKSVGEILDEELSGDRA------
SaCopG  1ea4 -----MKKRLTITLSESVLENLEKMAREMGLSK-SAMISVALENYKKGQek-------------------------------------------
EcAtaR  6gto -msavkkQRIDLRLTDDDKSMIEEAAAISNQSV-SQFMLNSASQRAAEVIEQHRRVilneeswtrvmdalsnppspgeklkraakrlqgm------
SoCopA  6iya ---mSTIKPVSVKLDADIKARVEHLAETRKRSS-HWMMREAIREYVEREEKREAL-----------------------------------------
P22 Arc 1arq mkgmskmPQFNLRWPREVLDLVRKVAEEN-GRSVNSEIYQRVMESFKKEGRIGA------------------------------------------
                    : : :       :   : :         :   :
```

**b.**

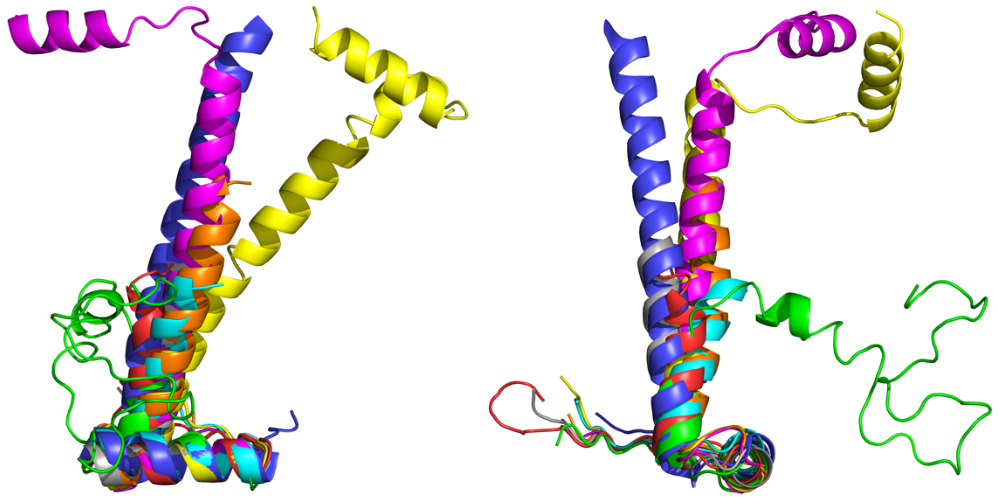

**Figure S2. A. Structure-based sequence alignment of *Vc*ParD2 and selected RHH proteins.** Residues that are not present in the model (because of disordered in the corresponding crystal structure) are shown in small letters. residues that have structurally equivalent residues in *Vc*ParD2 are shown in bold. Residues highlighted in green form the hydrophobic core of the *Vc*ParD2 dimer while those in cyan mediate the inter-dimer contacts in the higher order oligomer. Residues with side chains known to be involved in DNA recognition are highlighted in yellow. **B. Superposition** of the monomers of *Vc*ParD2 (orange), *Cc*ParD (magenta), *Mo*ParD3 (yellow) *RK2*ParD 2an7 (green), *Sa*CopG (cyan), *Ec*AtaR (blue), *So*CopA (grey) and P22 Arc (red).

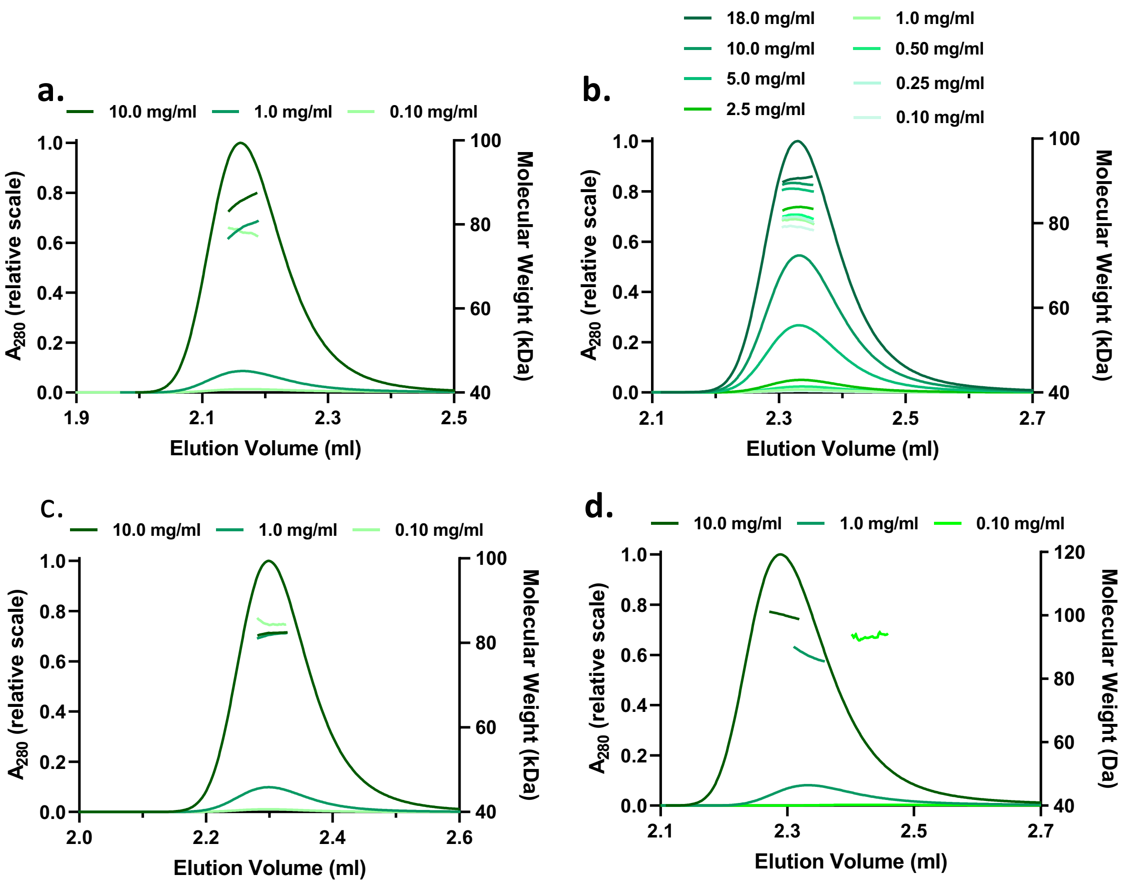


**Supplementary Figure S3. SEC-MALS**. Results of the SEC-MALS measurements showing the relative elution profiles and corresponding molecular weights of *Vc*ParD at concentrations varying from 18 mg/ml to 0.1 mg/ml. The experiments were performed in following buffer conditions: **a.** 20 mM Tris pH 8.0, 50 mM NaCl, 1mM TCEP; **b.** 20 mM Tris pH 8.0, 150 mM NaCl, 1mM TCEP; **c.** 20 mM Tris pH 8.0, 500 mM NaCl, 1mM TCEP; **d.** 50 mM NaAc pH 5.6, 150 mM NaCl, 1mM TCEP.

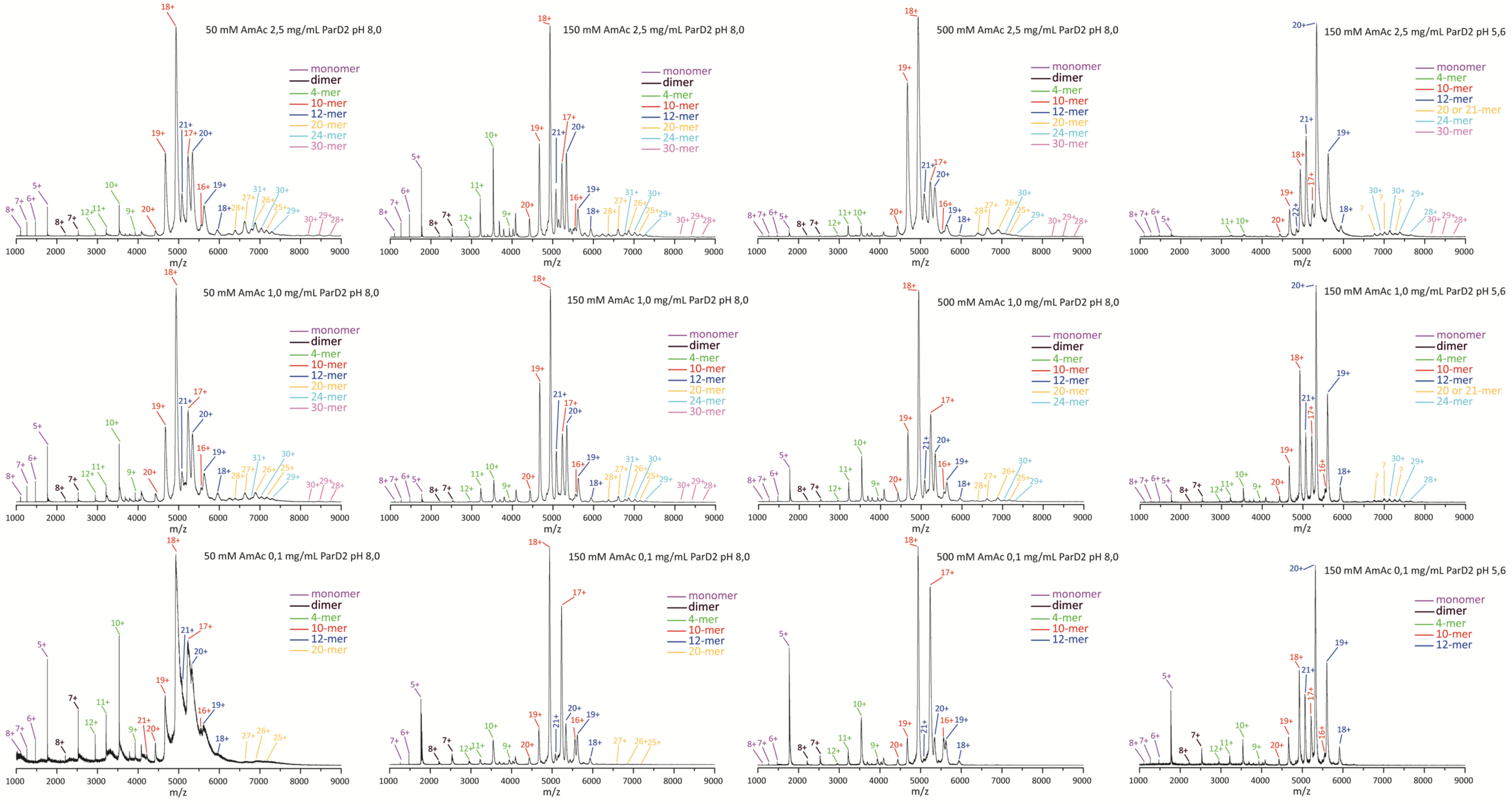


**Supplementary Figure S4. Native mass spectrometry.** Native MS spectra are shown for VcParD2 at different concetntrations, ionic strengts and pH. In all cases the dominant species are 10-mer and 12-mer althoughtheir relative abundance can vary.

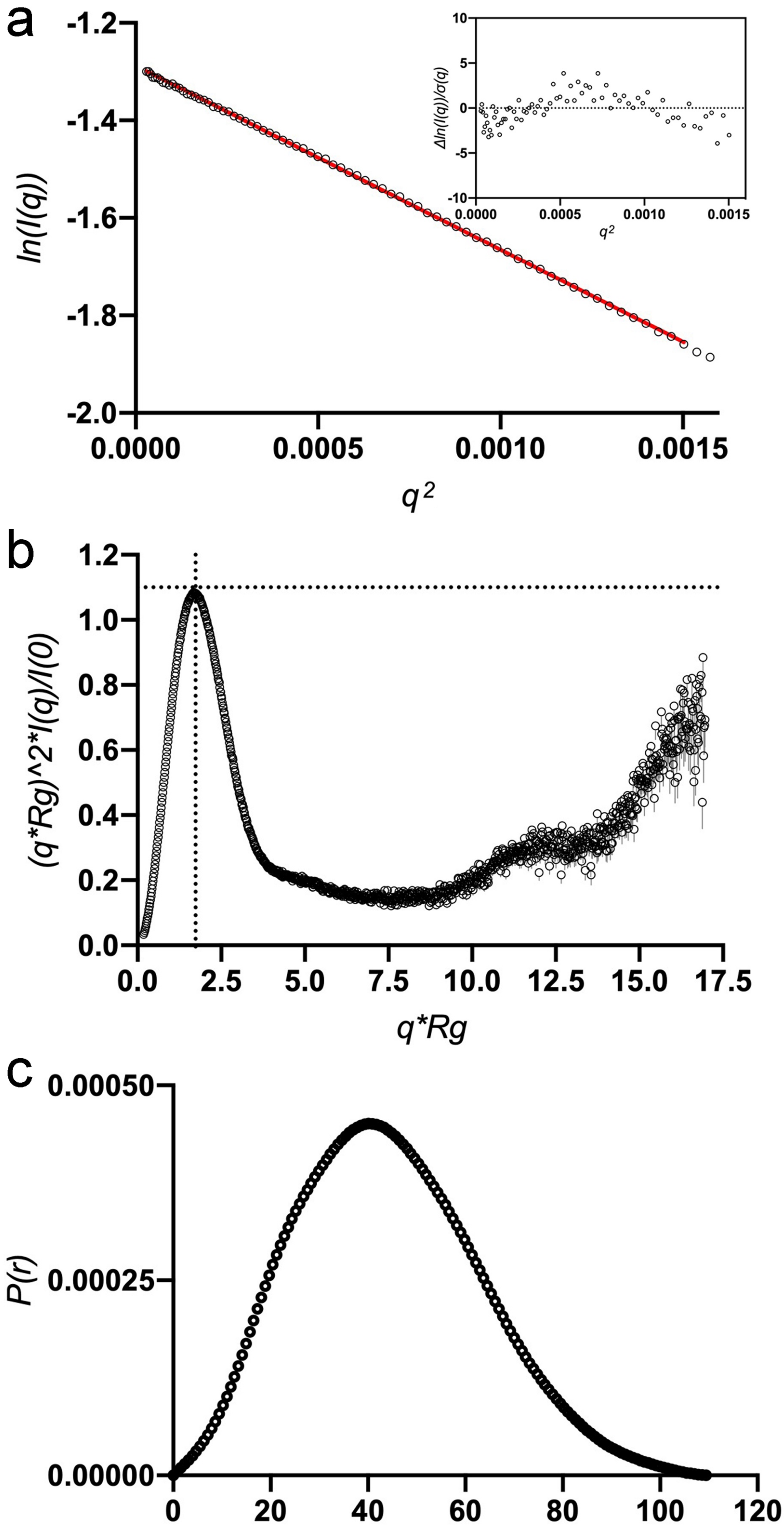


**Supplementary Figure S5. SAXS**. **A.** Guinier plot. The insert shows the residuals from the linear fit. **b**. Kratky plot. **c.** P(r) function.

### Supplementary Table I.

| concentration (mg/ml) | Ionic strength | Buffer | Technique | Molecular weight (kDa) |
|---|---|---|---|---|
| 0.15 | | | SEC | 170 |
| 12 | 150 mM NaCl | 20 mM Tris pH 8.0, 1 mM TCEP | SAXS (Porod) | 109 |
| 18 | 150 mM NaCl | 20 mM Tris pH 8.0, 1 mM TCEP | SEC-MALS | 91.0 ± 0.7% |
| 10 | 150 mM NaCl | 20 mM Tris pH 8.0, 1 mM TCEP | SEC-MALS | 89.1 ± 0.6% |
| 5 | 150 mM NaCl | 20 mM Tris pH 8.0, 1 mM TCEP | SEC-MALS | 87.7 ± 0.8% |
| 2.5 | 150 mM NaCl | 20 mM Tris pH 8.0, 1 mM TCEP | SEC-MALS | 83.4 ± 0.7% |
| 1 | 150 mM NaCl | 20 mM Tris pH 8.0, 1 mM TCEP | SEC-MALS | 79.9 ± 1.2% |
| 0.5 | 150 mM NaCl | 20 mM Tris pH 8.0, 1 mM TCEP | SEC-MALS | 80.4 ± 0.7% |
| 0.25 | 150 mM NaCl | 20 mM Tris pH 8.0, 1 mM TCEP | SEC-MALS | 80.6 ± 0.7% |
| 0.1 | 150 mM NaCl | 20 mM Tris pH 8.0, 1 mM TCEP | SEC-MALS | 78.5 ± 0.6% |
| 10 | 50 mM NaCl | 20 mM Tris pH 8.0, 1 mM TCEP | SEC-MALS | 85.0 ± 0.6% |
| 1 | 50 mM NaCl | 20 mM Tris pH 8.0, 1 mM TCEP | SEC-MALS | 79.2 ± 0.6% |
| 0.1 | 50 mM NaCl | 20 mM Tris pH 8.0, 1 mM TCEP | SEC-MALS | 77.9 ± 1.7% |
| 10 | 500 mM NaCl | 20 mM Tris pH 8.0, 1 mM TCEP | SEC-MALS | 81.9 ± 0.2% |
| 1 | 500 mM NaCl | 20 mM Tris pH 8.0, 1 mM TCEP | SEC-MALS | 81.6 ± 0.2% |
| 0.1 | 500 mM NaCl | 20 mM Tris pH 8.0, 1 mM TCEP | SEC-MALS | 84.8 ± 2.2% |
| 10 | 150 mM NaCl | 20 mM Na Acetate pH 5.6, 1 mM TCEP | SEC-MALS | 100.0 ± 0.6% |
| 1 | 150 mM NaCl | 20 mM Na Acetate pH 5.6, 1 mM TCEP | SEC-MALS | 87.6 ± 0.6% |
| 0.1 | 150 mM NaCl | 20 mM Na Acetate pH 5.6, 1 mM TCEP | SEC-MALS | 94.2 ± 1.9% |
